# Transcriptomic Dichotomy in Melanoma: Proliferative Cell Cycle Shield vs. Autoimmune-like Chromatin Activation in Anti-PD-1 Response

**DOI:** 10.1101/2025.05.09.653199

**Authors:** Basma Shabana, Ahmed Salem, Nouraldeen Ali Ramadan, Ahmed Mahmoud Khedr

**Affiliations:** Department of Dermatology, Menoufia University, Shebin El-Kom, El-Menoufia, Egypt; Bioinformatics Diploma Program, Nile University, Giza, Egypt; Faculty of Medicine, Tanta University, Tanta, Egypt; Neurology and Psychiatry Postgraduate Student, Faculty of Medicine, Ain Shams University, Cairo, Egypt; Cardiology and Vascular Medicine Postgraduate Student, Faculty of Medicine, Ain Shams University, Cairo, Egypt

**Keywords:** Melanoma immunotherapy, Anti-PD-1 resistance, Cell cycle shield, Chromatin remodeling, Transcriptomic signature, Biomarker discovery, Batch correction

## Abstract

Primary resistance to anti-PD-1 therapy in metastatic melanoma remains a clinical challenge. This study reanalyzed the GSE168204 dataset to elucidate molecular mechanisms of resistance and response, incorporating batch correction to address limitations in our prior preprint[6]. RNA-seq data from 25 melanoma biopsies (9 responders, 16 non-responders) were analyzed using DESeq2 with surrogate variable analysis[9,10]. We identified 3,247 differentially expressed genes, revealing a "cell cycle shield" signature in non-responders characterized by upregulation of CDK1, CCNB1, E2F1, and HSP90AA1 enriched for proliferation and DNA repair pathways, suggesting immune evasion through rapid tumor growth. Responders exhibited upregulation of EP300, CREBBP, FCGR2B, and histone genes enriched for chromatin organization and systemic lupus erythematosus pathways, indicating immune activation and autoimmune-like transcriptional programs. Notably, batch correction reversed the roles of EP300 and FCGR2B from non-responders to responders[6], highlighting their context-dependent functions in immune engagement. The "cell cycle shield" suggests targeting CDK1 or HSP90AA1 to overcome resistance[13,14], while the SLE signature may serve as a response biomarker reflecting immune activation states[7]. Validation in larger cohorts and experimental models is needed to translate these findings into personalized immunotherapy strategies.

## Introduction

Immune checkpoint inhibitors (ICIs) targeting programmed cell death protein 1 (PD-1) have revolutionized metastatic melanoma treatment, achieving durable responses in 40–50% of patients [1]. However, primary resistance remains a major challenge, with 40–60% of patients failing to respond to anti-PD-1 therapy [2]. While tumor mutational burden and PD-L1 expression partially predict response, they do not fully explain resistance mechanisms [3]. Emerging evidence suggests that tumor-intrinsic epigenetic and transcriptional programs, coupled with immune signaling, shape the tumor microenvironment to evade immune surveillance [4,5].Our previous preprint analyzed RNA-seq data from the GSE168204 dataset (n=25 melanoma biopsies) and proposed that epigenetic regulators, such as EP300, and immune inhibitory receptors, like FCGR2B, coordinate chromatin-driven immune evasion in non-responders to anti-PD-1 therapy [6]. However, potential batch effects in the original analysis may have confounded differential gene expression (DEG) profiles, prompting a reanalysis. In this updated study, we applied batch correction to the GSE168204 dataset, comparing responders (n=9) and non-responders (n=16) to anti-PD-1 therapy. Surprisingly, the reanalysis revealed that FCGR2B is upregulated in responders (logFC -1.39), potentially linked to immune activation, while EP300 shows only modest upregulation in responders (logFC -0.91), suggesting a less central role in resistance than previously hypothesized.The refined analysis identifies distinct transcriptional signatures: non-responders exhibit a “cell cycle shield” driven by proliferation and DNA repair pathways (e.g., CDK1, E2F1), while responders display immune-active profiles enriched for autoimmune pathways, such as systemic lupus erythematosus (SLE), which may correlate with immune-related adverse events (irAEs) [7]. These findings, supported by transcription factor and ligand-receptor interaction networks, provide a more reliable and nuanced understanding of resistance mechanisms. This study aims to elucidate the epigenetic and immune drivers of anti-PD-1 response in melanoma, offering insights into potential biomarkers and therapeutic targets to overcome resistance.

## Methods

### Data Acquisition and PreprocessingGene expression

Data were obtained from the GSE168204 dataset, comprising raw RNA-seq read counts from 25 melanoma biopsies (9 responders, 16 non-responders to anti-PD-1 therapy) deposited in the Gene Expression Omnibus (GEO) [8]. Raw counts were downloaded loaded into R (version 4.4.1) using the data.table package. Gene annotations were retrieved from the Human GRCh38.p13 annotation file, mapping gene IDs to symbols. Clinical metadata were extracted from the GSE168204 SOFT file. The response to immune checkpoint blockade therapy was categorized as responders (R) or non-responders (NR), and samples with missing response data were excluded. The count matrix and clinical metadata were aligned to include only common samples, resulting in a dataset of 25 samples with matching expression and clinical data.Genes with low expression (total counts < 1 across samples) were filtered out to reduce noise, retaining genes with sufficient read counts for downstream analysis. The filtered count matrix was used for differential expression analysis.

### Batch Correction and Differential Expression Analysis

To account for potential batch effects in the GSE168204 dataset, surrogate variable analysis (SVA) was performed using the sva package (version 3.50.0) [10]. A design matrix was constructed with response status (R vs. NR) as the primary variable, and a null model included only the intercept. The svaseq function identified significant surrogate variables (SVs), which were incorporated into the design matrix for differential expression analysis. Differential expression analysis was conducted using the DESeq2 package (version 1.42.0) [9]. A DESeqDataSet was created from the raw count matrix, including SVs and response status in the design formula. Results were extracted for the contrast of non-responders (NR) versus responders (R). Differentially expressed genes (DEGs) were defined as those with an adjusted p-value < 0.05 and |log2FC| > 0.5. DEGs were annotated with gene symbols using the GRCh38.p13 annotation file. Volcano plots were generated using the EnhancedVolcano package (version 1.20.0) to visualize DEGs, with upregulated genes (log2FC > 1.1, padj < 0.05) in red and downregulated genes (log2FC < -1.1, padj < 0.05) in blue. A heatmap of normalized counts for DEGs was created using the NMF package (version 0.27), with log2-transformed counts and sample annotations for response status.

### Pathway Enrichment AnalysisFunctional

enrichment analysis was performed separately for upregulated (log2FC > 0.5) and downregulated (log2FC < 0.5) DEGs in NR relative to R. Gene Ontology (GO) enrichment for Biological Process (BP), Molecular Function (MF), and Cellular Component (CC) was conducted using the clusterProfiler package (version 4.10.0) with the org.Hs.eg.db database (version 3.18.0). The enrichGO function was used with gene symbols as input, applying the Benjamini-Hochberg (BH) adjustment and a q-value cutoff of 0.05. KEGG pathway enrichment was conducted using enrichKEGG with gene IDs, organism code ‘hsa’, and BH adjustment. Reactome pathway enrichment was performed using the ReactomePA package (version 1.46.0). Enrichment results were visualized using bar plots and dot plots, showing the top 10–15 pathways per analysis.

### Transcription Factor (TF) Analysis

To identify transcription factors (TFs) regulating DEGs, the TRRUST v2 database was used, containing human TF-target interactions [11]. The TRRUST dataset was downloaded and loaded into R. A custom function mapped TF-target interactions to upregulated and downregulated DEG gene symbols. The top 10 TFs were identified based on the frequency of their target genes. Bar plots of TF frequencies were generated using base R, and TF-target networks were visualized using the tidygraph (version 1.3.0) and ggraph (version 2.1.0) packages. Nodes were annotated as TFs or target genes, with Fruchterman-Reingold layout and color-coded nodes.

### Receptor-TF and Ligand-Receptor Interaction Analysis

Receptor-TF interactions were retrieved from the OmniPath database using the OmnipathR package (version 3.2.0). All interactions for human (organism code 9606) were imported, and receptor-TF links were filtered to include source genes matching DEG symbols. The top 10 receptors by target count were visualized in bar plots using ggplot2 (version 3.5.0). Receptor-TF networks were constructed for the top 10 receptors and TFs using tidygraph and ggraph. Ligand-receptor interactions were also obtained from OmniPath (omnipath dataset). Interactions were filtered to include source genes in the DEG list, excluding self-interactions. Ligand-receptor networks were built for the top10 ligands and receptors, visualized using ggraph. Combined networks integrating ligand-receptor, receptor-TF, and TF-target edges were created, with nodes annotated as ligands, receptors, TFs, or target genes.

### Protein-Protein Interaction (PPI) Network Analysis

PPI networks were constructed using the STRINGdb package (version 2.14.0) [12] with STRING version 12 (score threshold: 400, species: human, 9606). Upregulated and downregulated DEGs were mapped to STRING IDs, and interactions were retrieved. PPI networks were built using igraph (version 2.0.3). Node attributes included gene symbols and log2FC, with betweenness and degree centrality calculated to identify hub genes. The top 20 hub genes were selected based on degree centrality. A combined edge and node attribute table was generated for Cytoscape visualization.

### Statistical Analysis and Visualization

All analyses were performed in R (version 4.4.1).Statistical significance for DEGs was determined using DESeq2’s Wald test with BH adjustment (padj < 0.05). Enrichment analyses used BH-adjusted q-values (< 0.05). Visualizations were generated using ggplot2, EnhancedVolcano, NMF, ggraph, and base R plotting functions.

## Results

### Differential Expression Analysis Reveals Distinct Signatures in Non-Responders and Responders

Differential expression analysis of the GSE168204 dataset (n=25 melanoma biopsies; 9 responders [R], 16 non-responders [NR] to anti-PD-1 therapy) identified 3,247 differentially expressed genes (DEGs) (supplemental table 1) after batch correction using surrogate variable analysis (SVA) (padj < 0.05, |log2FC| > 0.5). Of these, 1,678 genes were upregulated in NR (positive log2FC, higher in NR vs. R), and 1,569 were upregulated in R (negative log2FC, higher in R vs. NR). Volcano plots (Figure 1) and heatmaps (Figure 2) visualized the DEGs, highlighting distinct expression profiles between NR and R.In non-responders, upregulated genes were enriched for cancer/testis antigens (e.g., MAGEA10 [log2FC 8.29, padj 9.05e-07], MAGEA2 [log2FC 6.80, padj 2.09e-18]) and cell cycle-related genes (CDK1 [log2FC 1.26, padj 2.58e-02], CCNB1 [log2FC 1.16, padj 3.01e-02], PCNA [log2FC 1.10, padj 4.27e-03]) (Table 1). Notably, HDAC2 (log2FC 0.75, padj 3.48e-02) and CXCL8 (log2FC 2.36, padj 3.71e-02) indicated epigenetic modification and immune modulation, respectively. In contrast, responders showed upregulation of epigenetic modifiers (CHD5 [log2FC -4.92, padj 5.05e-05], KMT2A [log2FC -1.41, padj 2.11e-06], EP300 [log2FC -0.91, padj 9.49e-04]) and immune modulators (FCGR2B [log2FC -1.39, padj 3.34e-02], PDCD1LG2 [log2FC -0.99, padj 3.18e-02]). Histone genes, such as H1-3 (log2FC -5.87, padj 2.24e-09) and H3C11 (log2FC -5.01, padj 2.54e-04), were also significantly upregulated in R, suggesting active chromatin remodeling (Table 2).Contrary to our preprint findings [6], which suggested EP300 and FCGR2B as mediators of immune evasion in NR, batch correction revealed their upregulation in R. This shift indicates a role in immune activation rather than suppression, necessitating a reevaluation of their contribution to resistance mechanisms.

**Table 1:** Upregulated genes in NR.

| Symbol | log2FC | p-value | adj. p-value | Category |
| --- | --- | --- | --- | --- |
| MAGEA10 | 8.29 | 3.13e-09 | 9.05e-07 | Cancer/Testis Antigens |
| MAGEA2 | 6.80 | 5.76e-22 | 2.09e-18 | Cancer/Testis Antigens |
| MAGEA4 | 6.61 | 2.49e-04 | 5.54e-03 | Cancer/Testis Antigens |
| MAGEA9 | 5.44 | 1.51e-03 | 1.87e-02 | Cancer/Testis Antigens |
| MAGEA12 | 4.69 | 4.17e-07 | 5.61e-05 | Cancer/Testis Antigens |
| MAGEA3 | 3.53 | 1.11e-04 | 3.18e-03 | Cancer/Testis Antigens |
| MAGEA6 | 2.91 | 1.63e-03 | 1.96e-02 | Cancer/Testis Antigens |
| CDK5R1 | 2.18 | 2.23e-05 | 1.04e-03 | Cell Cycle Related |
| CDK1 | 1.26 | 2.45e-03 | 2.58e-02 | Cell Cycle Related |
| CCNB1 | 1.16 | 3.04e-03 | 3.01e-02 | Cell Cycle Related |
| PCNA | 1.10 | 1.72e-04 | 4.27e-03 | Cell Cycle Related |
| CDK20 | 1.05 | 2.26e-04 | 5.16e-03 | Cell Cycle Related |
| CDK5 | 0.82 | 1.88e-03 | 2.15e-02 | Cell Cycle Related |
| SMARCC1 | 0.68 | 4.42e-03 | 3.87e-02 | Cell Cycle Related |
| RBBP7 | 0.61 | 1.07e-04 | 3.09e-03 | Cell Cycle Related / Epigenetic Modifiers |
| NIFK | 0.61 | 3.71e-03 | 3.43e-02 | Cell Cycle Related |
| MSH2 | 0.54 | 1.51e-03 | 1.87e-02 | Cell Cycle Related |
| HDAC2 | 0.75 | 3.80e-03 | 3.48e-02 | Epigenetic Modifiers |
| H2AX | 0.87 | 4.82e-03 | 4.10e-02 | Histones |
| CXCL8 | 2.36 | 4.16e-03 | 3.71e-02 | Immune Modulators |
| PARP1 | 0.76 | 5.72e-03 | 4.60e-02 | Other |

**Table 2:** Upregulated genes in R.

| Symbol | log2FC | p-value | adj. p-value | Category |
| --- | --- | --- | --- | --- |
| CHD5 | -4.92 | 3.61e-07 | 5.05e-05 | Epigenetic Modifiers |
| KMT2A | -1.41 | 8.11e-09 | 2.11e-06 | Epigenetic Modifiers |
| KMT2D | -1.29 | 7.77e-06 | 4.83e-04 | Epigenetic Modifiers |
| EP300 | -0.91 | 1.95e-05 | 9.49e-04 | Epigenetic Modifiers |
| CHD2 | -0.85 | 2.19e-04 | 5.04e-03 | Epigenetic Modifiers |
| CREBBP | -0.77 | 9.49e-05 | 2.87e-03 | Epigenetic Modifiers |
| EZH1 | -0.76 | 1.51e-03 | 1.87e-02 | Epigenetic Modifiers |
| CHD9 | -0.73 | 4.23e-04 | 8.05e-03 | Epigenetic Modifiers |
| BANP | -0.69 | 1.43e-03 | 1.80e-02 | Epigenetic Modifiers |
| BICRAL | -0.67 | 1.86e-03 | 2.14e-02 | Epigenetic Modifiers |
| CHD8 | -0.56 | 1.83e-03 | 2.12e-02 | Epigenetic Modifiers |
| H1-3 | -5.87 | 3.25e-12 | 2.24e-09 | Histones |
| H3C11 | -5.01 | 3.16e-06 | 2.54e-04 | Histones |
| H4C13 | -4.69 | 5.31e-05 | 1.91e-03 | Histones |
| H3C2 | -4.27 | 1.89e-05 | 9.28e-04 | Histones |
| H2BC10 | -4.16 | 3.69e-05 | 1.47e-03 | Histones |
| H3-4 | -4.12 | 8.58e-04 | 1.29e-02 | Histones |
| H3C3 | -3.81 | 4.67e-04 | 8.54e-03 | Histones |
| H1-4 | -3.76 | 3.85e-09 | 1.04e-06 | Histones |
| H2BC14 | -3.53 | 3.29e-03 | 3.18e-02 | Histones |
| H4C2 | -3.37 | 5.95e-05 | 2.07e-03 | Histones |
| H2AC21 | -3.35 | 3.76e-05 | 1.49e-03 | Histones |
| H3C14 | -3.32 | 3.79e-07 | 5.22e-05 | Histones |
| H4C6 | -3.26 | 1.59e-04 | 4.05e-03 | Histones |
| H1-5 | -3.19 | 1.67e-04 | 4.18e-03 | Histones |
| H2AB2 | -3.19 | 1.57e-03 | 1.91e-02 | Histones |
| H3C15 | -3.17 | 1.62e-06 | 1.59e-04 | Histones |
| H3C1 | -3.17 | 1.90e-05 | 9.30e-04 | Histones |
| H4C1 | -3.16 | 6.85e-05 | 2.29e-03 | Histones |
| H2BC8 | -3.14 | 9.45e-04 | 1.37e-02 | Histones |
| H2BC3 | -2.96 | 3.35e-03 | 3.21e-02 | Histones |
| H2AC8 | -2.79 | 6.03e-04 | 1.01e-02 | Histones |
| H2AC14 | -2.67 | 8.69e-04 | 1.29e-02 | Histones |
| H3C6 | -2.60 | 1.51e-04 | 3.96e-03 | Histones |
| H2BC7 | -2.15 | 5.75e-03 | 4.62e-02 | Histones |
| H2AC12 | -2.08 | 1.47e-03 | 1.84e-02 | Histones |
| H4C3 | -2.04 | 1.31e-04 | 3.58e-03 | Histones |
| H4C5 | -2.02 | 2.33e-04 | 5.29e-03 | Histones |
| H2BC26 | -1.73 | 1.58e-03 | 1.93e-02 | Histones |
| H4C16 | -1.56 | 6.77e-04 | 1.08e-02 | Histones |
| H2BC18 | -1.31 | 9.77e-04 | 1.40e-02 | Histones |
| FCGR2B | -1.39 | 3.57e-03 | 3.34e-02 | Immune Modulators |
| PDCD1LG2 | -0.99 | 3.30e-03 | 3.18e-02 | Immune Modulators |
| PARP2 | -1.52 | 1.89e-05 | 9.28e-04 | Other |
| MLH3 | -1.03 | 8.66e-04 | 1.29e-02 | Other |
| CDKL5 | -0.94 | 8.49e-06 | 5.16e-04 | Other |

### Functional Pathway Enrichment Highlights Proliferation in NR and Immune Activation in R

Pathway enrichment analysis was performed on upregulated DEGs in NR and R using Gene Ontology (GO), KEGG, and Reactome databases (Figures 3–8). In NR(figures 3-5), upregulated genes were significantly enriched for ribosome biogenesis (e.g., ribonucleoprotein complex biogenesis, rRNA processing), cell cycle and DNA replication (e.g., chromosome segregation, DNA strand elongation), DNA repair (e.g., double-strand break repair), and mitochondrial function (e.g., oxidative phosphorylation) (Table 3). These pathways support a hyperproliferative, metabolically adapted phenotype in NR, termed the “cell cycle shield,” which likely contributes to immune evasion and resistance to anti-PD-1 therapy (Table 3). In responders(figures 6-8), upregulated genes were enriched for chromatin organization (e.g., nucleosome assembly, protein-DNA complex organization), immune pathways (e.g., antigen receptor-mediated signaling, neutrophil extracellular trap formation), and autoimmune-related pathways, notably systemic lupus erythematosus (SLE) (KEGG: hsa05322). Additional enrichments included calcium signaling, cell adhesion, and hematopoietic differentiation (e.g., RUNX1, NOTCH pathways), reflecting an open chromatin state and active immune engagement (Table 4). The SLE pathway reflects activation of autoimmunity-related gene programs in R. While such patterns have been linked to immune-related toxicities in other contexts, our dataset does not allow direct clinical correlation.

**Table 3:**
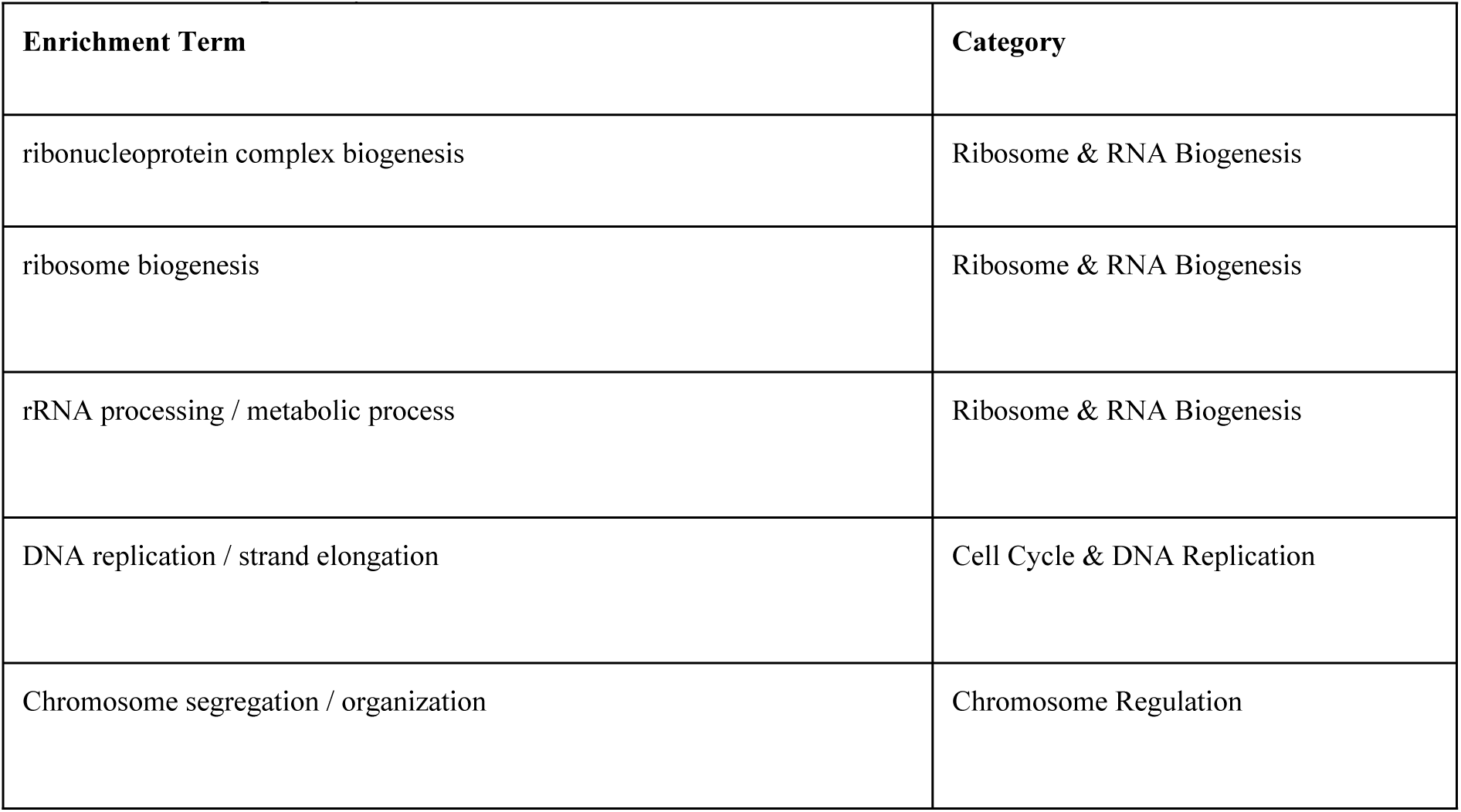
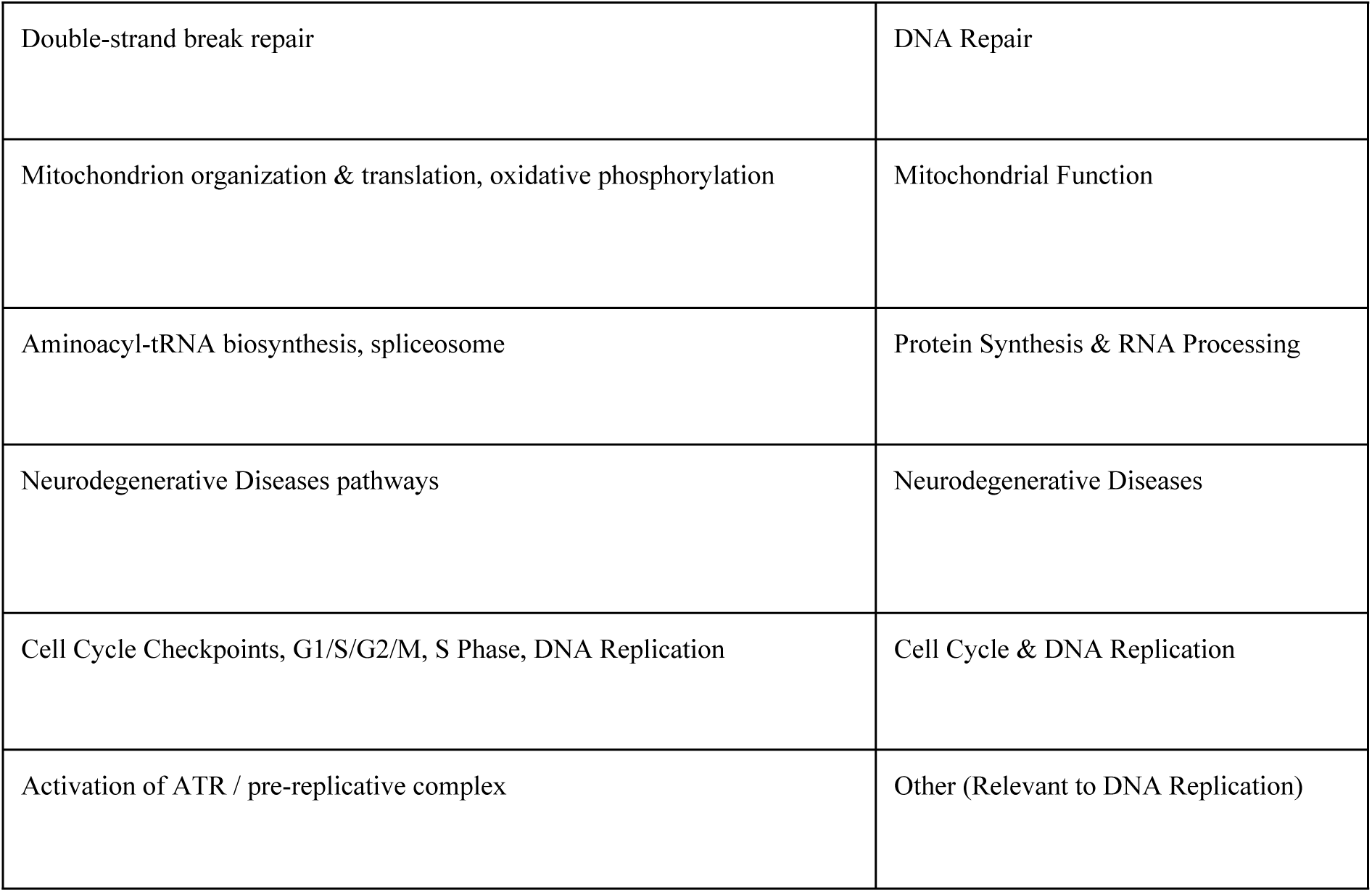
Enchired pathways in NR.

**Table 4:** Enchired pathways in R.

| Enrichment Term | Category |
| --- | --- |
| nucleosome organization / assembly, protein-DNA complex organization/assembly | Chromatin Organization |
| calcium ion transmembrane/transport | Ion Transport & Signaling |
| regulation of small GTPase mediated signal transduction, signal transduction | Signal Transduction |
| cell-substrate adhesion | Cell Adhesion |
| antigen receptor-mediated signaling pathway, Neutrophil extracellular trap formation, Toxoplasmosis | Immune Pathways |
| Systemic lupus erythematosus (KEGG SLE), Transcriptional misregulation in cancer | Autoimmune & Immune Pathways, Transcriptional Regulation |
| HDACs deacetylate histones, DNA methylation, chromatin modifications | Epigenetic Regulation |
| RUNX1/megakaryocyte, granulopoiesis, NOTCH, androgen receptor signaling | Hematopoiesis/Differentiation |
| RNA Polymerase I Promoter Opening | Transcriptional Regulation |
| Reproduction, Renin secretion, Sensory pathways (taste), Oxytocin, Hypertrophic cardiomyopathy | Various |

A side-by-side comparison (Table 5) underscores the contrast: NR tumors are dominated by proliferation and biosynthetic pathways, while R tumors exhibit epigenetic and immune activation, facilitating anti-tumor immunity.

**Table 5:** side-by-side comparison of enchired pathways between NR and R.

| Functional Class | Main Enrichment in NR | Main Enrichment in R |
| --- | --- | --- |
| Cell Cycle & DNA Replication | Strongly upregulated (cell cycle, DNA rep., repair) | Minimal |
| Ribosome & RNA Biogenesis | Upregulated | Not prominent |
| Mitochondrial Function | Upregulated | Not prominent |
| Chromatin Organization | Not prominent | Upregulated - nucleosome organization |
| Epigenetic Regulation | — | Active (HDACs, DNA methylation, chromatin modifications) |
| Immune Pathways | Minor | Strong - SLE, NETs, antigen receptor signaling |
| Signal Transduction | Not much | Diverse (GTPases, calcium, oxytocin) |
| Cell Adhesion | Minimal | Upregulated |
| Neurodegenerative diseases | Upregulated (feature of aggressive tumors/metabolism) | — |
| Hematopoietic Differentiation | — | Active (RUNX1, granulopoiesis, NOTCH) |
| Autoimmune Pathways | — | Upregulated (SLE) |

Transcription Factor Networks Reveal Context-Dependent RegulationTranscription factor (TF) analysis using the TRRUST v2 database [11] identified key TFs regulating DEGs in NR and R (Figure 9-10). In NR(figure 9), top TFs included E2F1, SP1, RELA, NFKB1, and TP53, associated with cell cycle progression, inflammation, and DNA damage response (Table 6). Related genes (AATF, ABL1, ATM) further supported stress response and transcriptional control, reinforcing the proliferative phenotype. In R(figure 10), SP1, NFKB1, and RELA were upregulated, while FOXA1, EBF1, and FLI1 were prominent, linked to immune differentiation and chromatin remodeling (Table 6). The opposite regulation of shared TFs (SP1, NFKB1, RELA) suggesting their context-dependent roles: promoting proliferation in NR and supporting immune activation in R. Receptor and Ligand Interactions Support Distinct Tumor Microenvironments. Receptor-TF(Table 9) and ligand-receptor interactions were analyzed using OmniPath (Figures 11–16). In NR(figure 11,13), E2F1 (983 targets) and HDAC2 (438 targets) were major receptor hubs, regulating cell cycle inhibitors (CDKN2A, CDKN1A) and DNA repair genes (BRCA1) (Table 7). Ligand-receptor pairs(figure 14), such as MIR17HG-CDKN1A and SRC-EGFR, underscored oncogenic signaling and proliferation (Table 10). In R(figure 12,15), EP300 (2119 targets) and FOXA1 (1130 targets) were dominant, regulating tumor suppressors (TP53) and immune genes (SMAD2, CTNNB1). Ligand-receptor interactions(figure 16), including EP300-TP53 and CREBBP-CTNNB1, supported chromatin remodeling and immune activation (Table 8,10). These networks highlight a proliferative, epigenetically repressed state in NR versus an open chromatin, immune-active state in R.

**Table 6:** TFs and related genes for both NR and R.

| Group | TF/Genes | Functional Categories |
| --- | --- | --- |
| **Top TFs in NR** | SP1, E2F1, RELA, NFKB1, TP53, EGR1, MYC, MYCN, HDAC1, YY1 | Cell cycle regulation (E2F1, MYC), Inflammation/immune signaling (RELA, NFKB1, EGR1), Tumor suppressor (TP53), Chromatin modification (HDAC1), General transcription regulation (SP1, YY1) |
| **Top genes related to NR TFs** | AATF, ABL1, AHR, AR, ARID3A, ATF3, ATF4, ATF6, ATF7, ATM | DNA damage response (ATM, AATF), Transcription factors (ATFs), Hormone receptor (AR), Chromatin remodeling (ARID3A) |
| **Top TFs in R** | SP1, NFKB1, RELA, EGR1, YY1, SP3, ETS1, HIF1A, JUN, PPARG | Transcription regulation (SP1, YY1, SP3), Immune and stress response (NFKB1, RELA, EGR1, JUN, HIF1A), Metabolic regulation (PPARG), TF involved in differentiation (ETS1) |
| **Top genes related to R TFs** | AES, AR, ARID3A, ARNTL, ATF2, ATM, BATF, BRCA1, BTF3, CDX2 | Transcription factors (AES, ARID3A, ARNTL, ATF2, BATF, BTF3, CDX2), DNA repair (BRCA1, ATM), Hormone receptor (AR) |

**Table 7:** top receptors in NR.

| Group | Receptor (Source Gene) | Number of Target Genes | Category / Functional Role |
| --- | --- | --- | --- |
| **NR (Non-Responders)** | E2F1 | 983 | Cell cycle TF, proliferation driver |
|  | HDAC2 | 438 | Epigenetic modifier, chromatin remodeling |
|  | MAFF | 236 | Stress response TF, oxidative stress response |
|  | SMARCC1 | 82 | Chromatin remodeler (SWI/SNF complex) |
|  | TRIM28 | 77 | Transcriptional co-repressor, epigenetics |
|  | ELK1 | 41 | MAPK pathway TF, proliferation and stress |
|  | SOX10 | 18 | Neural crest development, melanoma marker |
|  | RARB | 6 | Retinoic acid receptor, differentiation |
|  | KLF1 | 4 | Developmental TF |
|  | SOHLH1 | 4 | Transcription factor, germ cell development |

**Table 8:** top receptors in R.

| Group | Receptor (Source Gene) | Number of Target Genes | Category / Functional Role |
| --- | --- | --- | --- |
| **R**<br>**(Responders)** | EP300 | 2119 | Histone acetyltransferase, co-activator |
|  | FOXA1 | 1130 | Pioneer TF, chromatin remodeling |
|  | EBF1 | 698 | B cell TF, differentiation |
|  | ZEB1 | 310 | EMT regulator, transcriptional repressor |
|  | CHD2 | 243 | Chromatin remodeler (CHD family) |
|  | FOXP1 | 176 | Immune TF, T cell regulation |
|  | BDP1 | 93 | RNA Pol III transcription factor |
|  | POU5F1 | 85 | Stemness and pluripotency TF |
|  | MYCN | 67 | Oncogenic TF, cell proliferation |
|  | FLI1 | 17 | ETS family TF, hematopoiesis/immune |

**Table 9:**
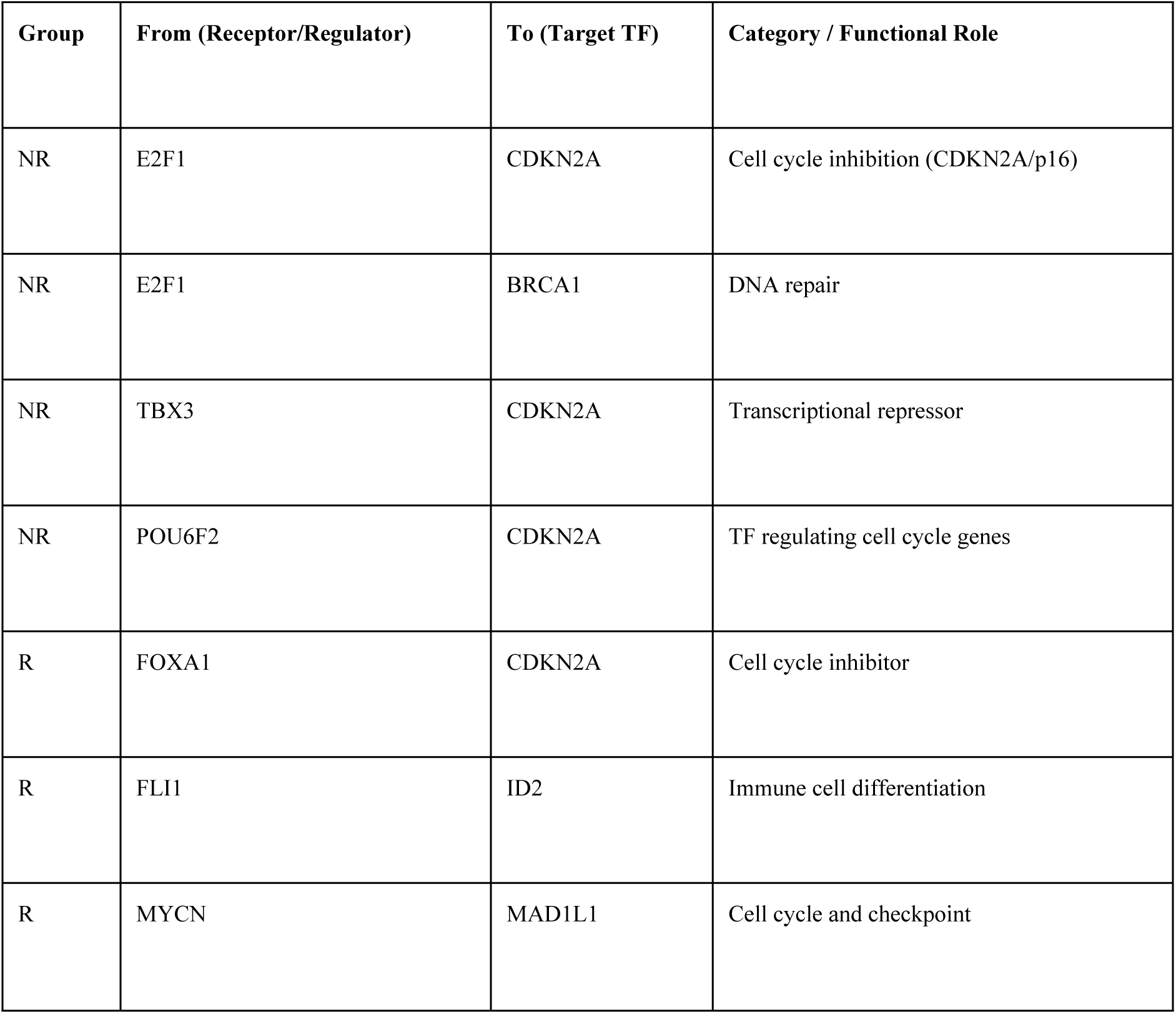
recptors to TFs.

| Group | From (Receptor/Regulator) | To (Target TF) | Category / Functional Role |
| --- | --- | --- | --- |
| NR | E2F1 | CDKN2A | Cell cycle inhibition (CDKN2A/p16) |
| NR | E2F1 | BRCA1 | DNA repair |
| NR | TBX3 | CDKN2A | Transcriptional repressor |
| NR | POU6F2 | CDKN2A | TF regulating cell cycle genes |
| R | FOXA1 | CDKN2A | Cell cycle inhibitor |
| R | FLI1 | ID2 | Immune cell differentiation |
| R | MYCN | MAD1L1 | Cell cycle and checkpoint |

**Table 10:** top ligand to receptors.

| Group | From (Ligand) | To (Receptor/Target Gene) | Functional Role / Interpretation |
| --- | --- | --- | --- |
| NR | MIR17HG | CDKN1A | miRNA cluster regulating cell cycle inhibitor p21, promoting proliferation in NR. |
| NR | E2F1 | CDKN1A, CDK1 | Autoregulation of cell cycle components supporting unchecked proliferation. |
| NR | PARP1 | HIF1A, ATM | DNA repair and hypoxia response pathway activation, enhancing cell survival. |
| NR | SRC | EGFR | Oncogenic signaling driving proliferation and resistance mechanisms. |
| NR | TRIM28 | STAT3 | Transcriptional repressor influencing immune evasion and tumor growth via STAT3 signaling. |
| NR | PCNA | DNMT1 | DNA replication and epigenetic modification coupling, promoting tumor cell survival. |
| R | EP300 | TP53, H3C12, CTNNB1, SMAD2 | Activates tumor suppressor TP53 and chromatin remodeling/immune signaling pathways favoring response. |
| R | CREBBP | CTNNB1, H3C12, SMAD2 | Chromatin modification and signal transduction supporting immune activation and tumor control. |
| R | KMT2A | H3C12 | Histone methyltransferase linked to active chromatin states in responders. |
| R | CASP8 | AR | Apoptosis-related signaling possibly modulating tumor suppression and immune signaling. |

### Protein-Protein Interaction Networks Identify Key Hub Genes

Protein-protein interaction (PPI) networks were constructed using STRING for upregulated DEGs in NR and R. In NR, hub genes included HSP90AA1, CDK1, CCNB1, CDC20, and PCNA, associated with cell cycle progression, DNA replication, and protein homeostasis (Table 11). These hubs reflect a tumor phenotype resistant to immunotherapy through rapid proliferation and stress tolerance. In R, hub genes such as TNF, EP300, CREBBP, SETD2, CD8A, and CD74 indicated immune signaling, epigenetic activation, and antigen presentation (Table 12). Histone genes (H3C13, H4C6) further suggested an open chromatin state conducive to immune activity.

**Table 11:**
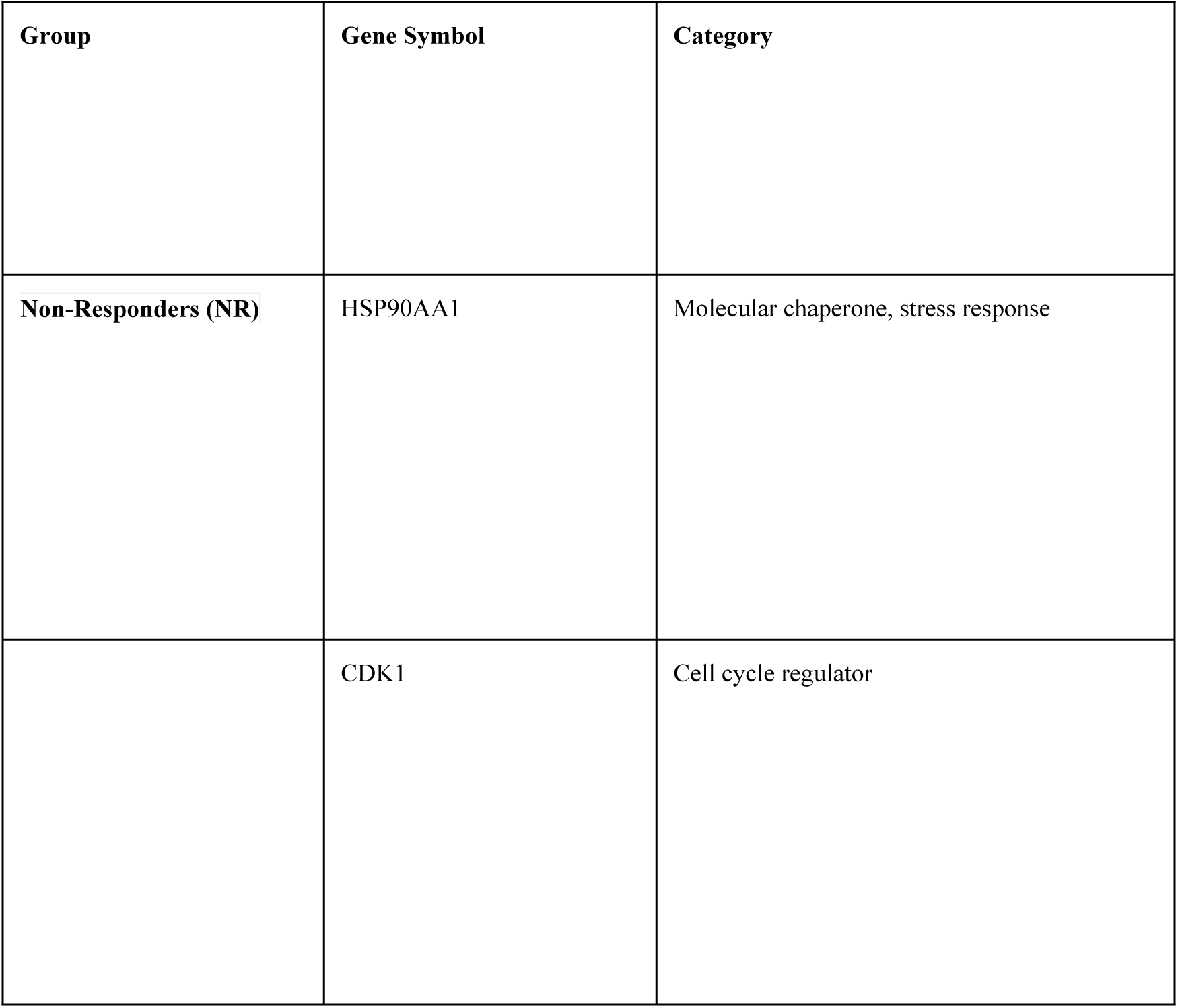
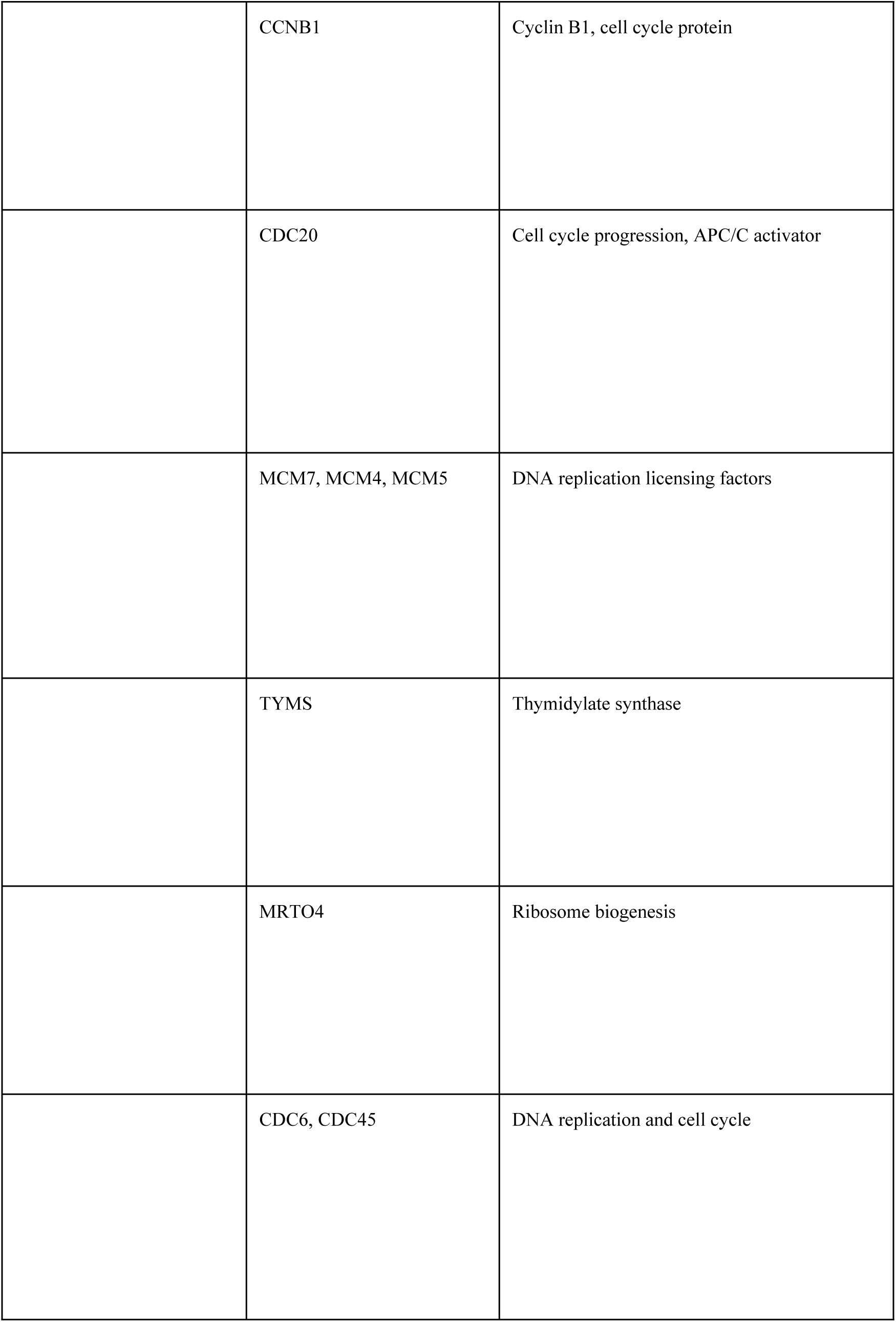
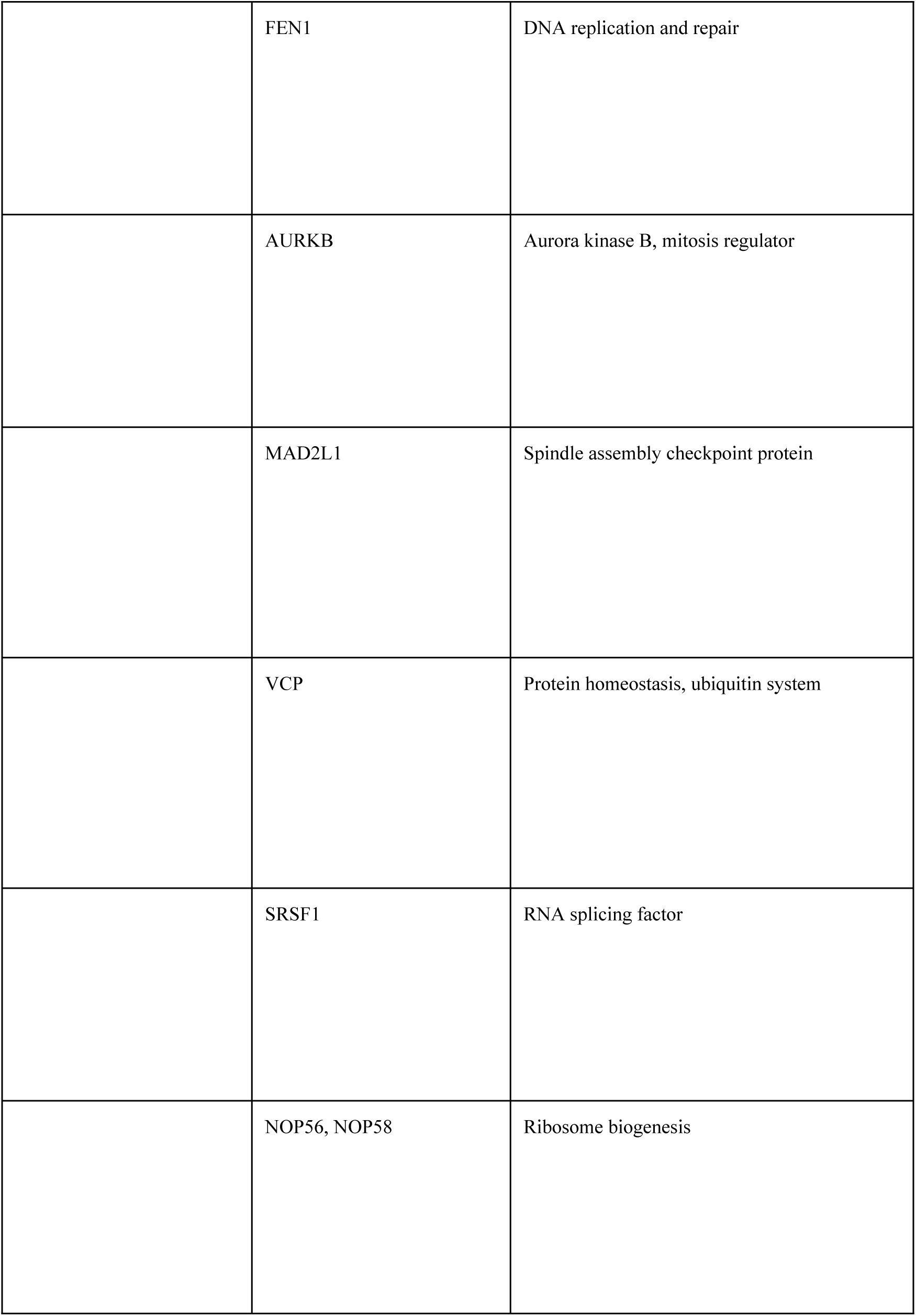
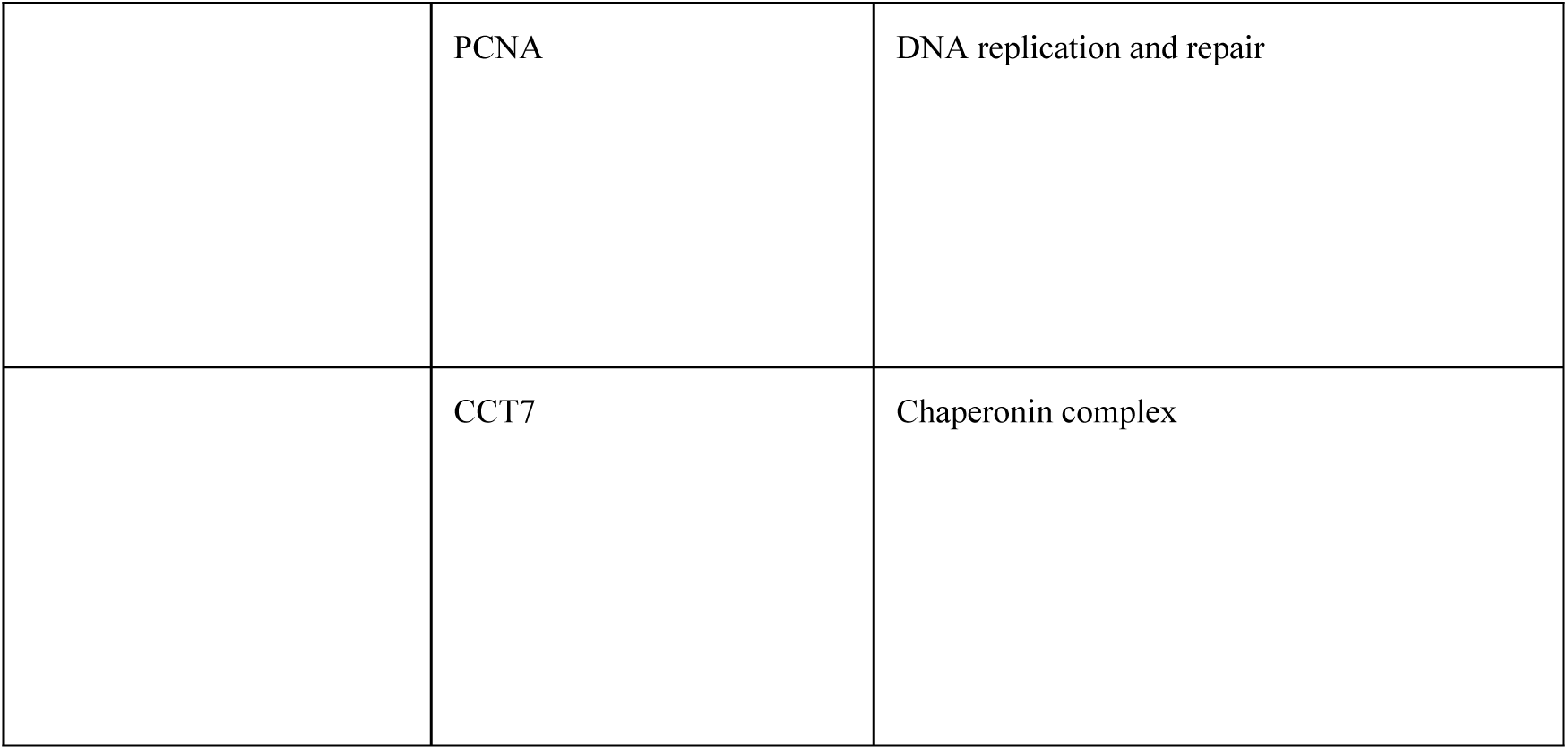
top hub protiens in NR.

**Table 12:** top hub protiens in R.

| Group | Gene Symbol | Category |
| --- | --- | --- |
| **Responders**<br>**(R)** | TNF | Immune signaling, cytokine |
|  | EP300 | Histone acetyltransferase (HAT),<br>transcription coactivator |
|  | H3C13, H4C6, H3-4, H2BC3, H2AC8, H2BC14,<br>H2AC14, H3C1, H3C3, H3C6 | Core histone proteins |
|  | CD8A | Immune cell marker, T cell coreceptor |
|  | CREBBP | Histone acetyltransferase, transcriptional<br>coactivator |
|  | SETD2 | Histone methyltransferase |
|  | KMT2A, KMT2C, KMT2D | Histone methyltransferases |
|  | CD74 | MHC class II invariant chain |
|  | CACNA1C | Calcium channel subunit |

### Key Findings and Implications

The reanalysis with batch correction revealed critical insights:

1. Reversal of EP300 and FCGR2B Roles: Unlike the preprint [6], EP300 and FCGR2B are upregulated in R, suggesting roles in immune activation and chromatin remodeling rather than immune evasion in NR.
2. Cell Cycle Shield in NR: Upregulation of CDK1, CCNB1, E2F1, and DNA repair genes in NR supports a proliferative, resistant phenotype, potentially targetable with cell cycle inhibitors [13,14].
3. Immuno-related Signature in R: The enrichment of SLE and immune pathways in R, coupled with TNF and CD8A, aligns with clinical observations of irAEs in responders [7], suggesting a biomarker for response, potentially reflecting immune activation states.
4. Epigenetic Dichotomy: NR tumors exhibit epigenetic repression (HDAC2, TRIM28), while R tumors show active chromatin remodeling (EP300, CREBBP, KMT2A), facilitating immune engagement.

These findings redefine the molecular landscape of anti-PD-1 resistance and response in melanoma, highlighting actionable pathways and potential biomarkers.

## Discussion

Our reanalysis of the GSE168204 dataset, incorporating batch correction via surrogate variable analysis (SVA), has significantly refined our understanding of the molecular mechanisms underlying primary resistance to anti-PD-1 therapy in metastatic melanoma. The identification of 3,247 differentially expressed genes (DEGs) revealed distinct transcriptional profiles between non-responders (NR) and responders (R), overturning key findings from our preprint [6]. Notably, the reversal of EP300 and FCGR2B expression from NR to R, coupled with the identification of a proliferation-driven “cell cycle shield” in NR and an autoimmune-linked signature in R, provides novel insights into resistance mechanisms and potential therapeutic targets.

The most striking finding is the unexpected upregulation of EP300 (log2FC -0.91, padj 9.49e-04) and FCGR2B (log2FC -1.39, padj 3.34e-02) in responders, contrasting with our preprint’s hypothesis that these genes drive immune evasion in NR [6]. EP300, a histone acetyltransferase, promotes open chromatin states and transcriptional activation of immune-related genes. Its upregulation in R, alongside other epigenetic modifiers like CREBBP, KMT2A, and SETD2, suggests an active chromatin landscape that facilitates anti-tumor immunity. Similarly, FCGR2B, an inhibitory Fcγ receptor, Its upregulation in R may reflect a context-dependent role in modulating immune responses, potentially enhancing effector cell activity or contributing to immune-related adverse events (irAEs) [7]. This reversal underscores the importance of batch correction in mitigating technical artifacts, as uncorrected batch effects in the preprint likely led to false positives. These findings align with studies showing that epigenetic activation and immune modulation are critical for immunotherapy response [4, 5].

In non-responders, the upregulation of cell cycle and DNA repair genes (CDK1, CCNB1, PCNA, E2F1) and cancer/testis antigens (MAGEA10, MAGEA2) supports a hyperproliferative phenotype, termed the “cell cycle shield.” This signature, enriched for ribosome biogenesis, DNA replication, and mitochondrial function, enables tumor cells to evade immune surveillance by prioritizing growth and survival. E2F1 and HDAC2, major regulatory hubs in NR, drive cell cycle progression and epigenetic repression, respectively, consistent with mechanisms of immune evasion. The prominence of HSP90AA1 as a hub gene further suggests stress tolerance and protein homeostasis as key resistance mechanisms [17]. These findings align with reports linking cell cycle dysregulation to immunotherapy resistance, suggesting that targeting CDK1 or HSP90 with inhibitors could sensitize NR tumors to anti-PD-1 therapy [13,14].

In responders, the enrichment of immune pathways, particularly systemic lupus erythematosus (SLE) (KEGG: hsa05322), alongside TNF, CD8A, and CD74, indicates robust immune activation and antigen presentation. The immuno-related signature is particularly intriguing, as it may reflect an autoimmune-like state in responders, which in other contexts have been linked to immune-related toxicities; however, we cannot confirm this association here. This is supported by the upregulation of histone genes (H1-3, H3C11) and epigenetic regulators (EP300, CREBBP), which facilitate open chromatin and active transcription of immune genes. The receptor-TF network, driven by EP300 and FOXA1, further supports tumor suppressor activation (TP53) and immune signaling (SMAD2), creating a tumor microenvironment conducive to immunotherapy response. These findings suggest that epigenetic therapies, such as HDAC inhibitors, could enhance response rates by mimicking the open chromatin state of responders [16].

The context-dependent regulation of shared transcription factors (SP1, NFKB1, RELA) between NR and R highlights their dual roles. In NR, these TFs promote proliferation and inflammation, while in R, their upregulation may suppress tumor growth and enhance immune activation. This dichotomy underscores the complexity of transcriptional networks in melanoma and the need for context-specific therapeutic strategies.

In conclusion, this reanalysis redefines the molecular basis of anti-PD-1 resistance in melanoma. The “cell cycle shield” in NR suggests targeting proliferation pathways (CDK1, HSP90AA1) to overcome resistance, while the autoimmuno-relted and epigenetic signatures in R highlight potential biomarkers for response and irAEs. These findings pave the way for personalized therapeutic strategies, combining cell cycle inhibitors or epigenetic modulators with immunotherapy to improve outcomes in melanoma patients.

## Conclusion

This reanalysis of the GSE168204 dataset, incorporating batch correction via surrogate variable analysis, has elucidated distinct molecular signatures underlying primary resistance to anti-PD-1 therapy in metastatic melanoma. The identification of 3,247 differentially expressed genes revealed a proliferation-driven “cell cycle shield” in non-responders, characterized by upregulation of CDK1, CCNB1, E2F1, and HSP90AA1, which supports immune evasion through rapid tumor growth and stress tolerance. In contrast, responders exhibited an epigenetically active, immune-engaged phenotype, marked by upregulation of EP300, CREBBP, FCGR2B, and a systemic lupus erythematosus SLE-autoimmuno-related signature, representing activation of autoimmune-like transcriptional programs, without direct clinical correlation in this dataset. The reversal of EP300 and FCGR2B expression from non-responders in our preprint [6] to responders in this study underscores their roles in chromatin remodeling and immune activation, highlighting the critical impact of batch correction in refining molecular insights.

These findings suggest actionable therapeutic strategies, including targeting cell cycle regulators (CDK1, HSP90AA1) with inhibitors to overcome resistance in NR [13,14] and leveraging epigenetic modulators (e.g., HDAC inhibitors) to enhance immune activation in R [16]. The autoimmuno-related signature in responders offers a potential biomarker for predicting response, though its relationship to irAEs requires validation in cohorts with toxicity data[7]. Future studies should validate key genes (CDK1, EP300, FCGR2B) using experimental approaches (e.g., qPCR, CRISPR) and larger cohorts, such as TCGA melanoma, to confirm their roles and explore their therapeutic potential. By delineating the molecular dichotomy between resistance and response, this study provides a foundation for personalized immunotherapy strategies to improve outcomes in metastatic melanoma.

## Limitations and Future Directions

While this reanalysis of the GSE168204 dataset provides novel insights into anti-PD-1 therapy resistance in metastatic melanoma, several limitations must be acknowledged. First, the small sample size (n=25; 9 responders, 16 non-responders) and uneven group distribution may limit statistical power and generalizability, potentially introducing bias in differential expression analysis despite batch correction with surrogate variable analysis (SVA) [10]. Second, the study relies on bioinformatics analysis without experimental validation, which restricts the confirmation of key findings such as the roles of EP300, FCGR2B, CDK1, and the autoimmuno-related signature. Third, the dataset lacks detailed clinical metadata (e.g., specific irAE profiles), limiting the ability to correlate molecular findings with clinical outcomes like immune-related adverse events (irAEs) [7]. fourth, batch effects, while mitigated, may still influence subtle gene expression differences, particularly for low-abundance transcripts. Finally tumer heterogenesity which can resovle by single cell analysis[15].

Future studies should address these limitations through experimental validation of key genes (CDK1, EP300, FCGR2B) using techniques such as quantitative PCR (qPCR), CRISPR-based gene editing, or immunohistochemistry in independent melanoma cohorts. Validation in larger datasets could confirm the generalizability of the “cell cycle shield” in non-responders and the immuno-related signature in responders. Integrating single-cell RNA sequencing could elucidate cell-type-specific contributions to resistance and response, particularly for immune modulators like FCGR2B. Clinical studies correlating the immuo-related signature with irAE incidence and severity are warranted to establish its utility as a biomarker [7]. Therapeutically, combining cell cycle inhibitors (e.g., targeting CDK1 or HSP90AA1) or epigenetic modulators (e.g., HDAC inhibitors) with anti-PD-1 therapy should be explored in preclinical models to overcome resistance [13,14,16]. These efforts will refine our understanding of resistance mechanisms and facilitate personalized immunotherapy strategies for melanoma patients.

## Data Availability

The GSE168204 dataset is publicly available at the Gene Expression Omnibus (https://www.ncbi.nlm.nih.gov/geo/query/acc.cgi?acc=GSE168204).

---

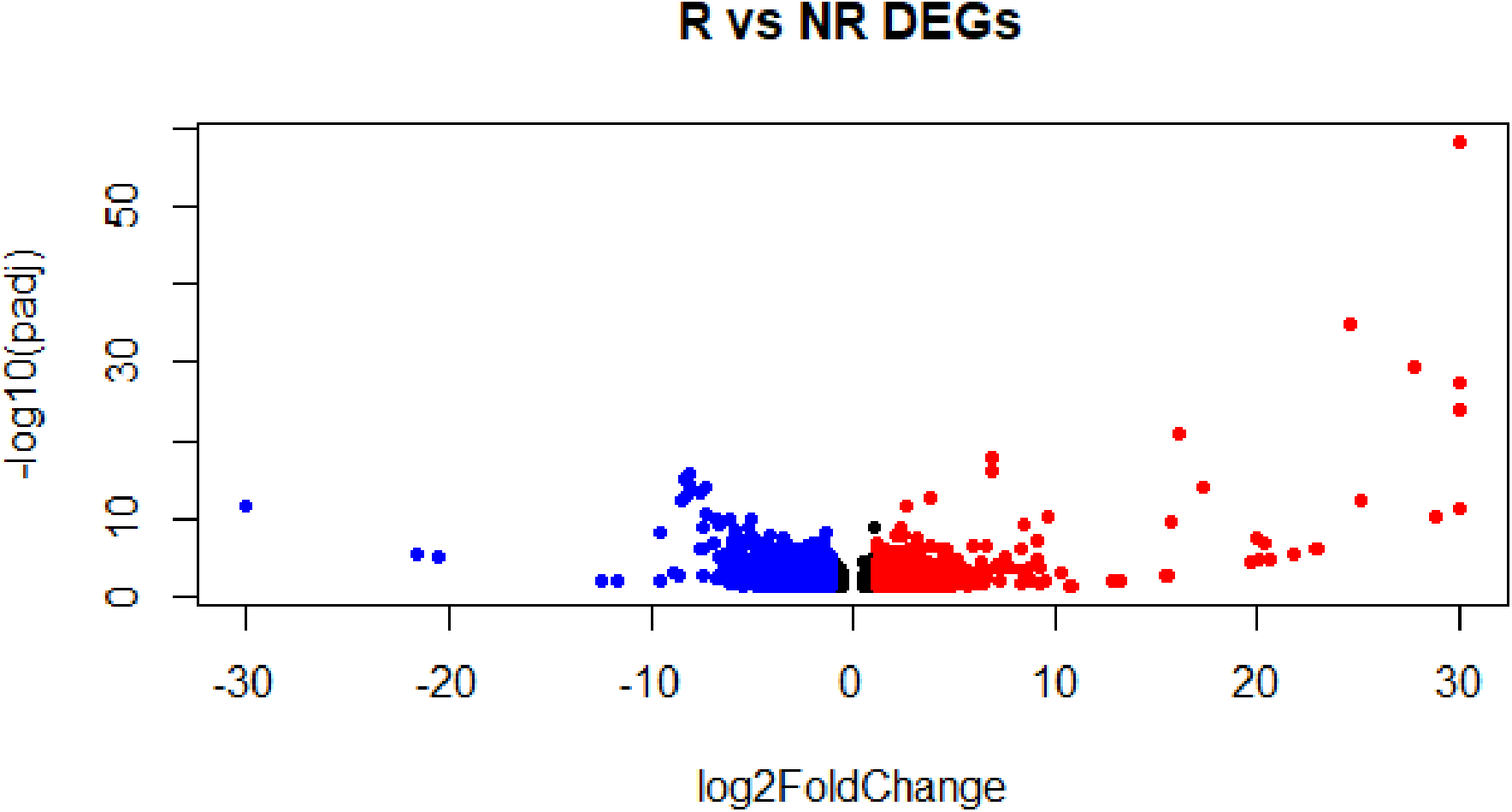

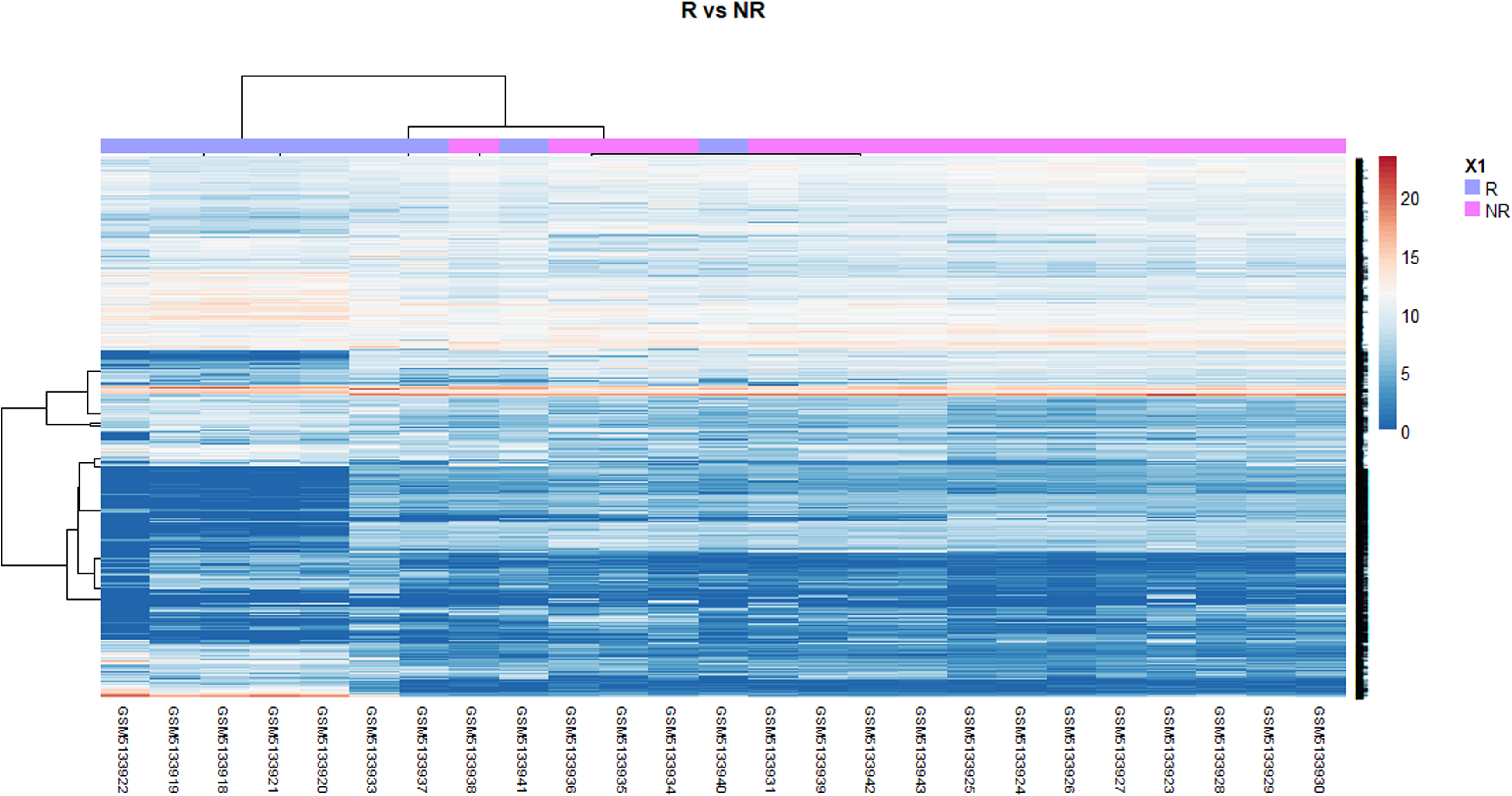

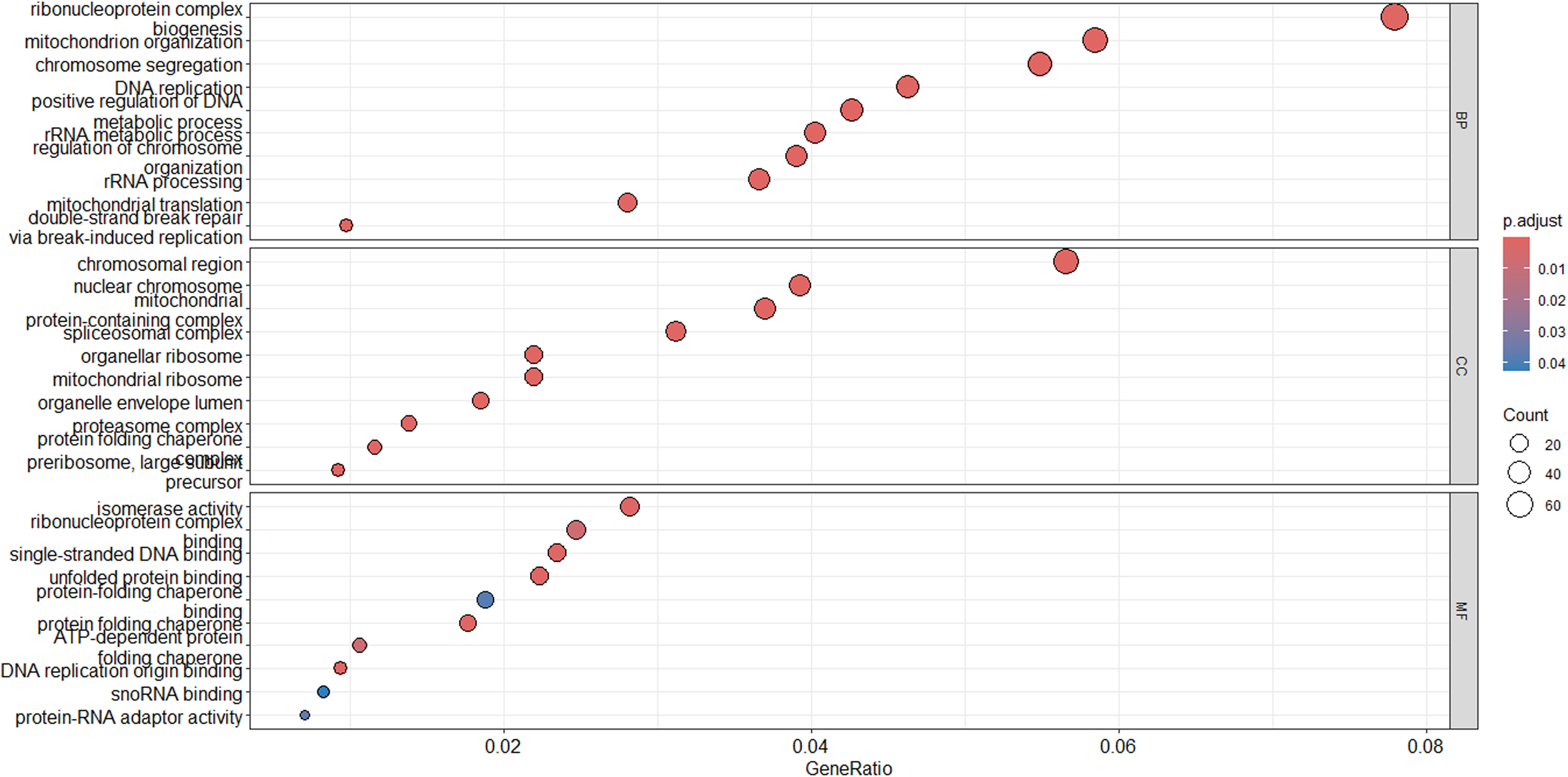

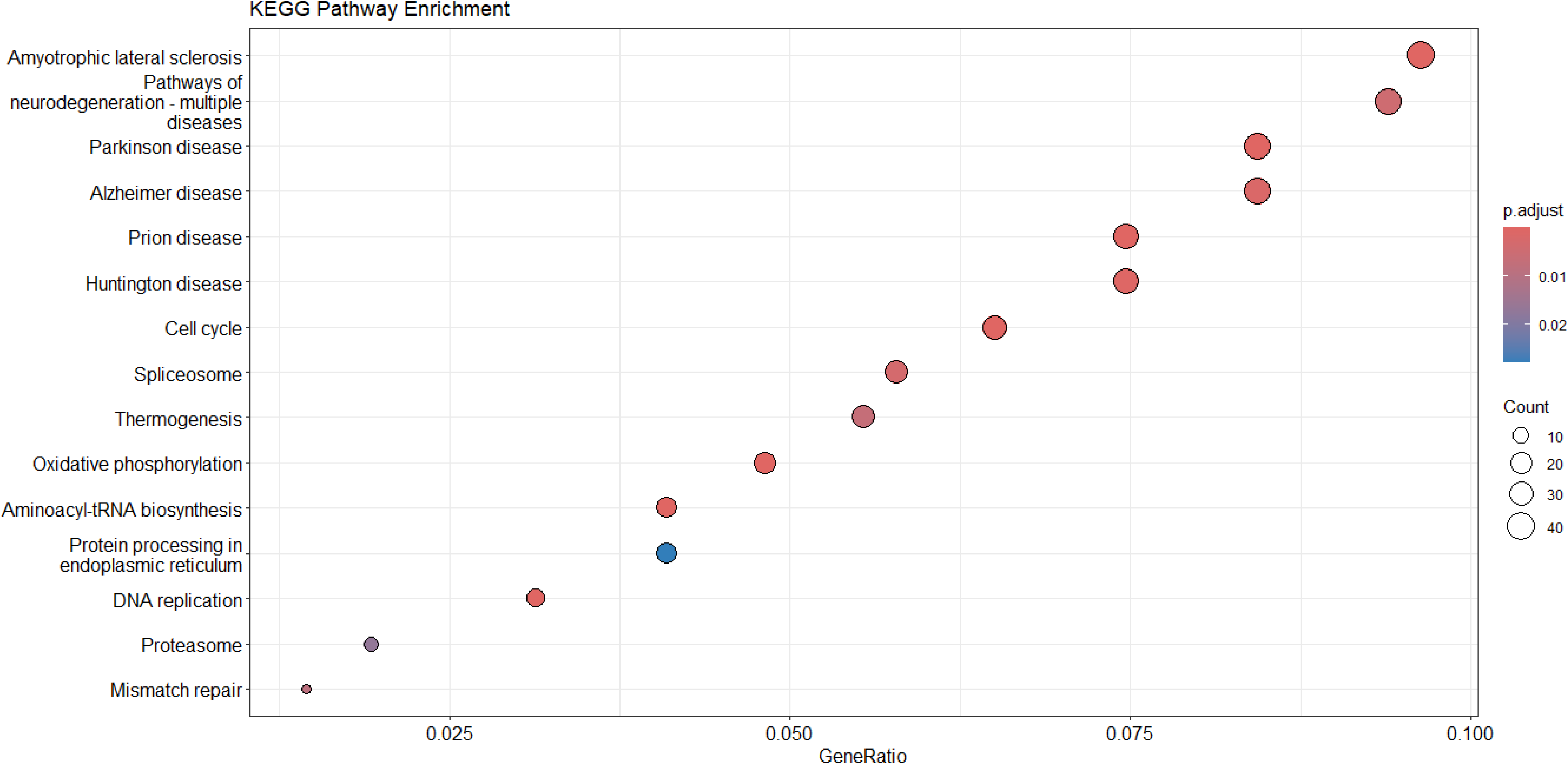

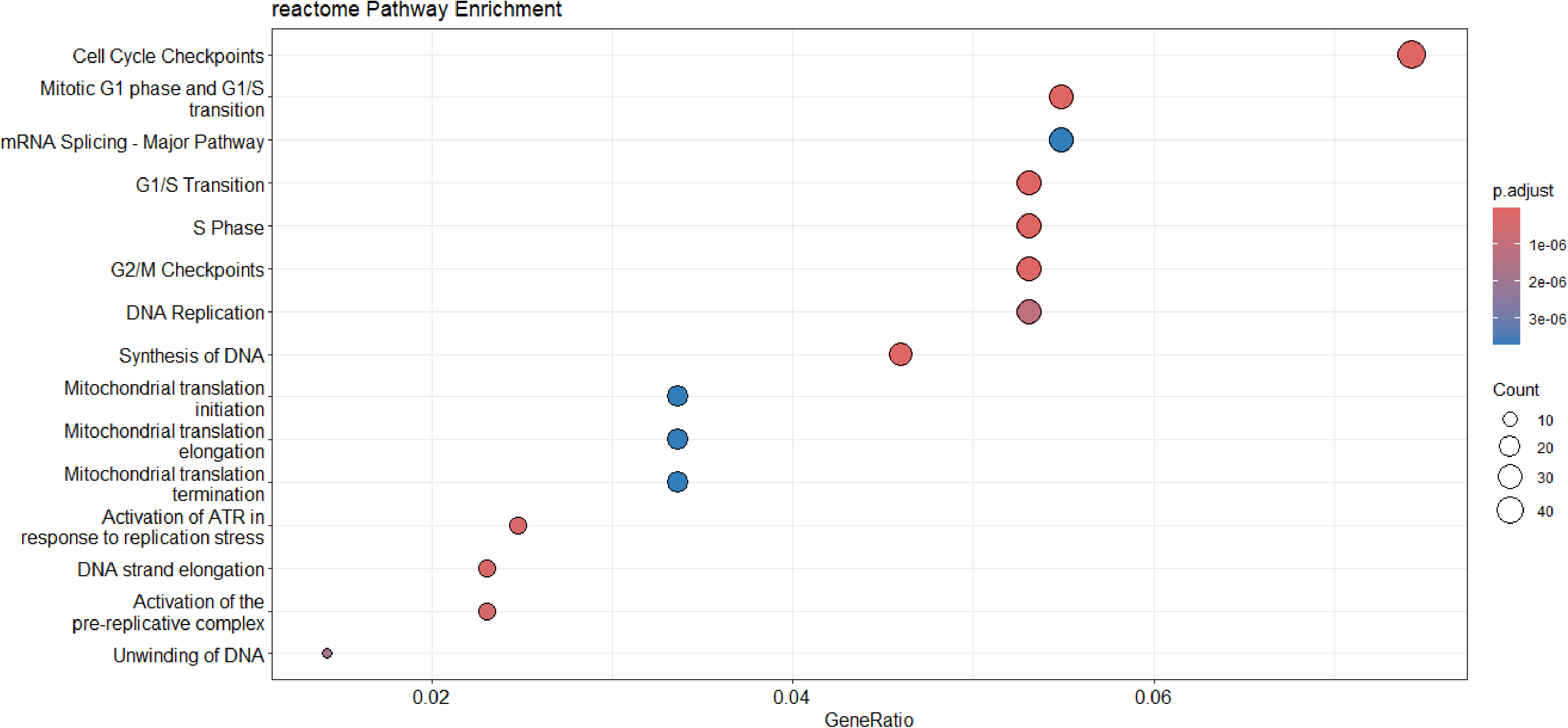

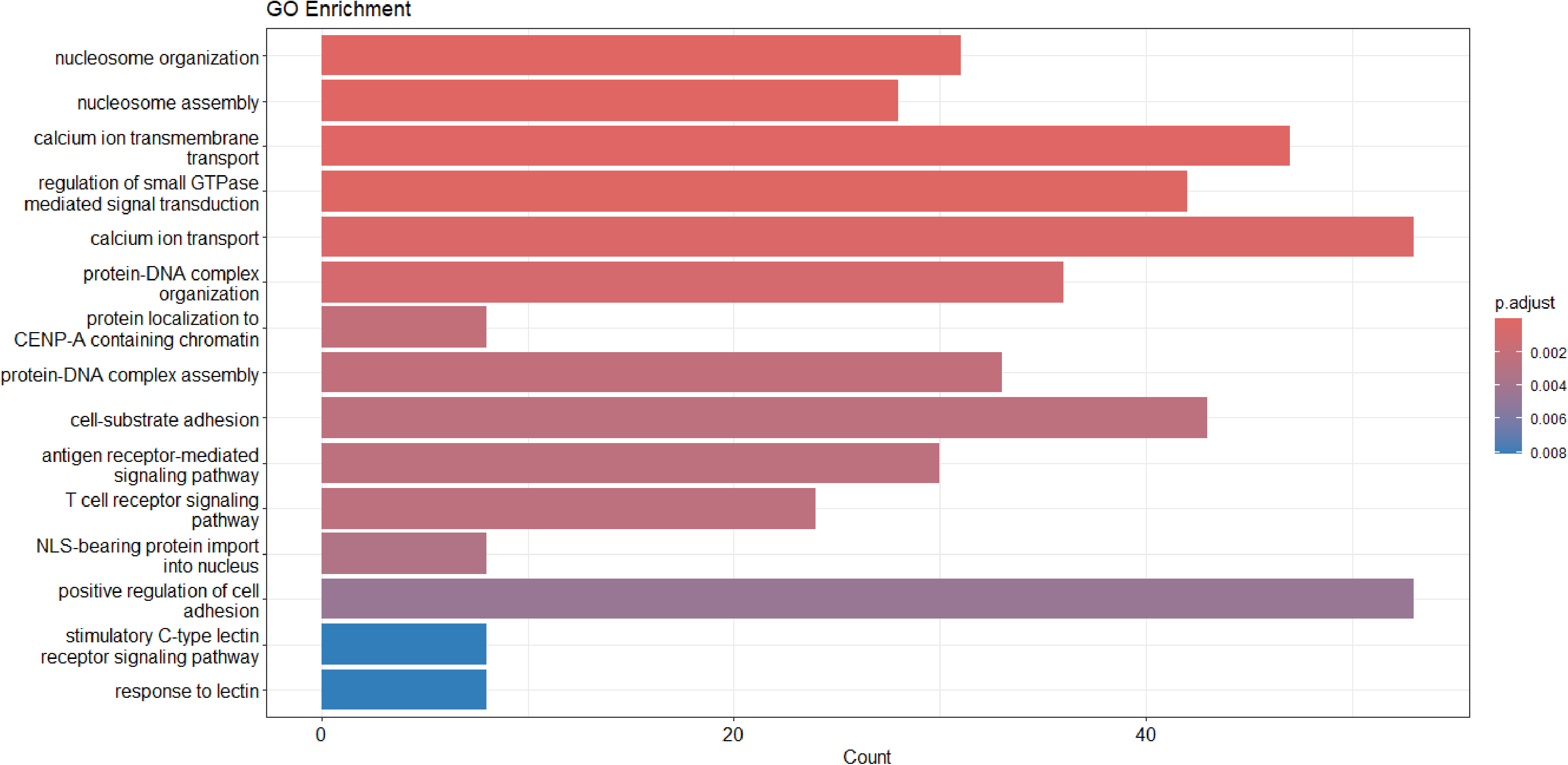

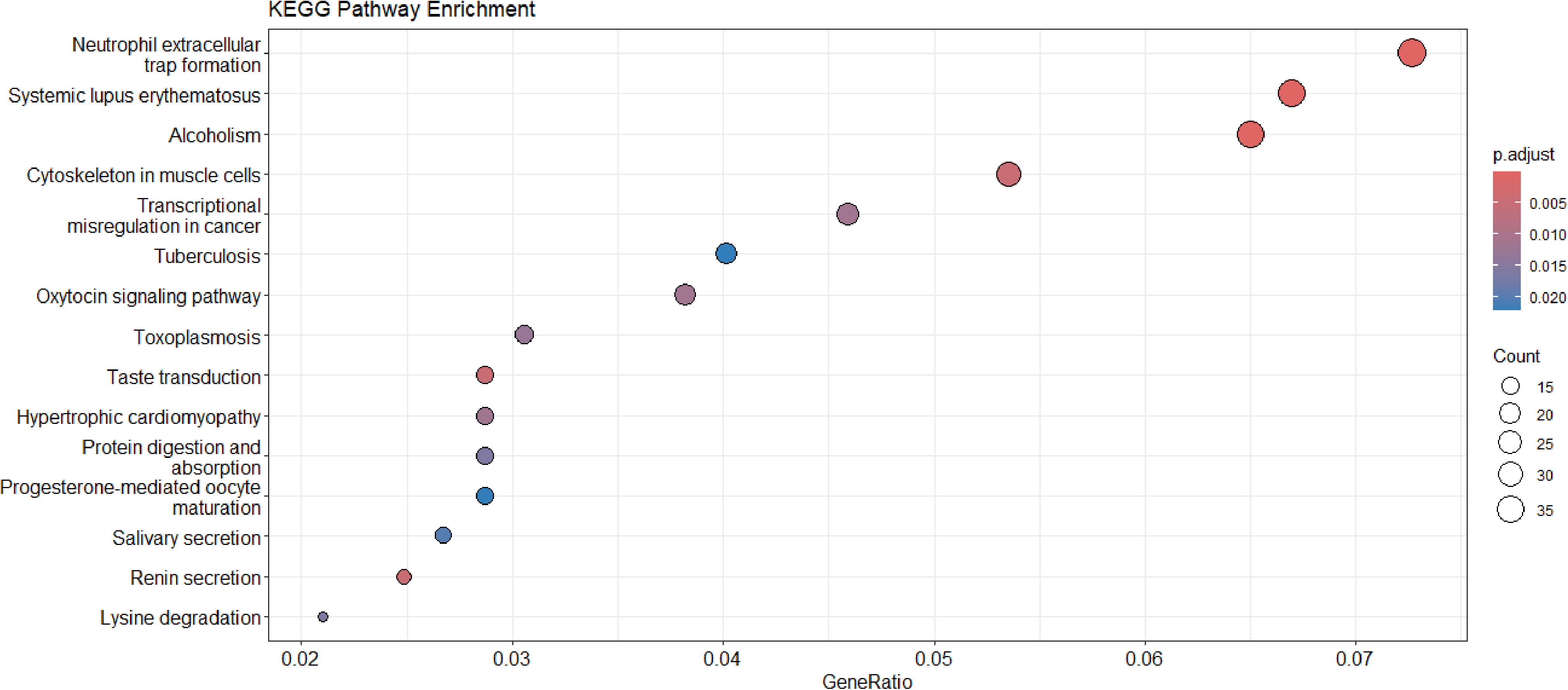

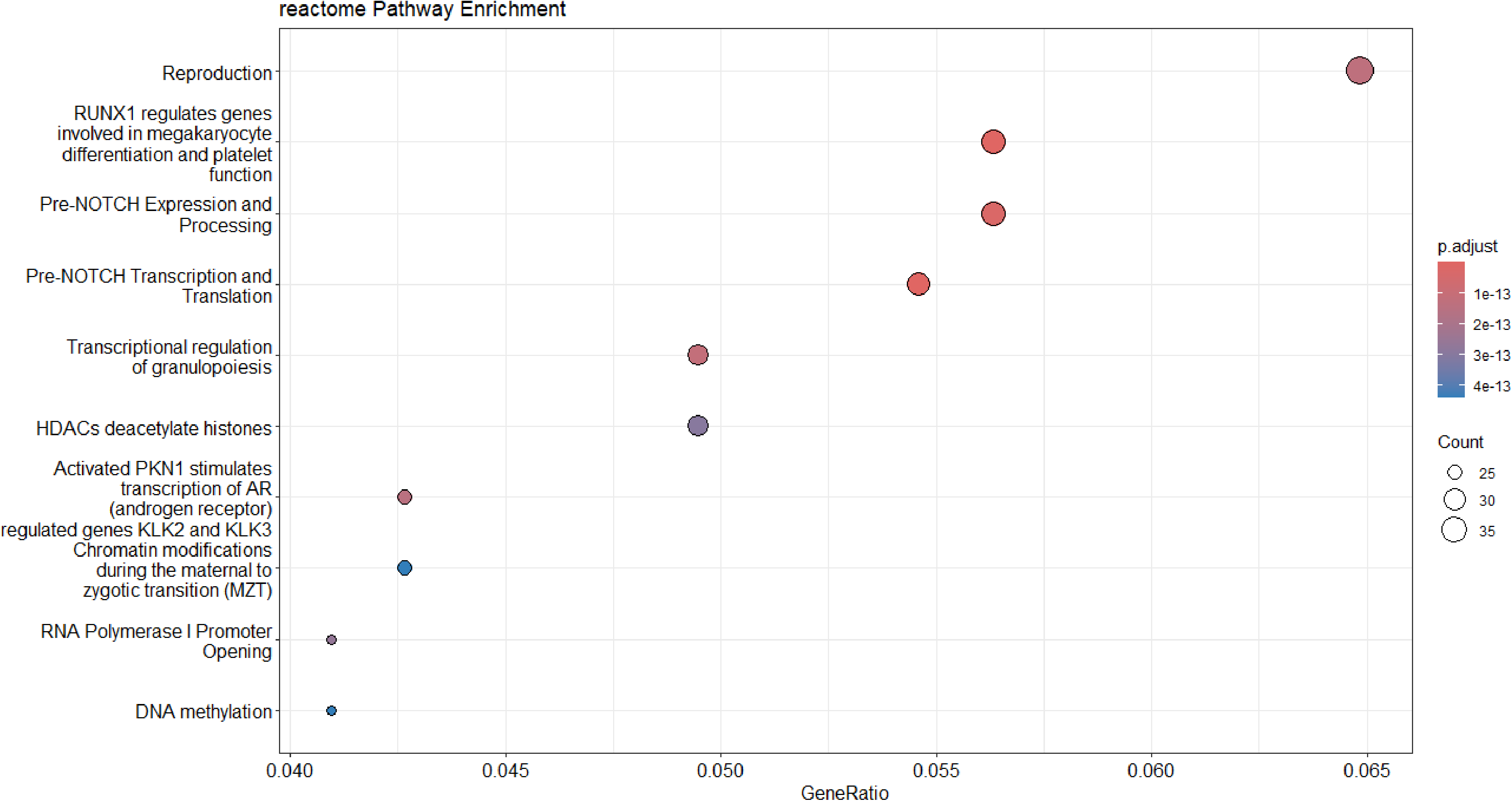

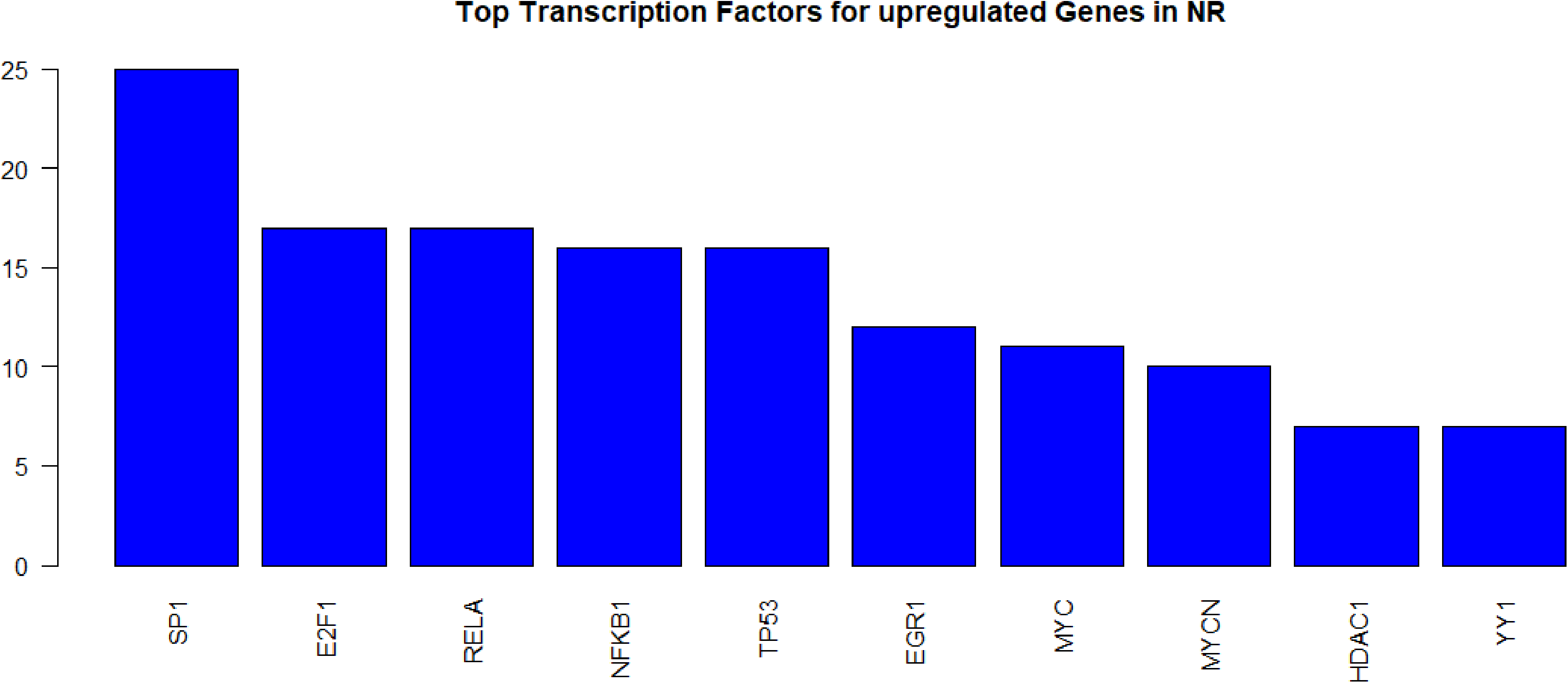

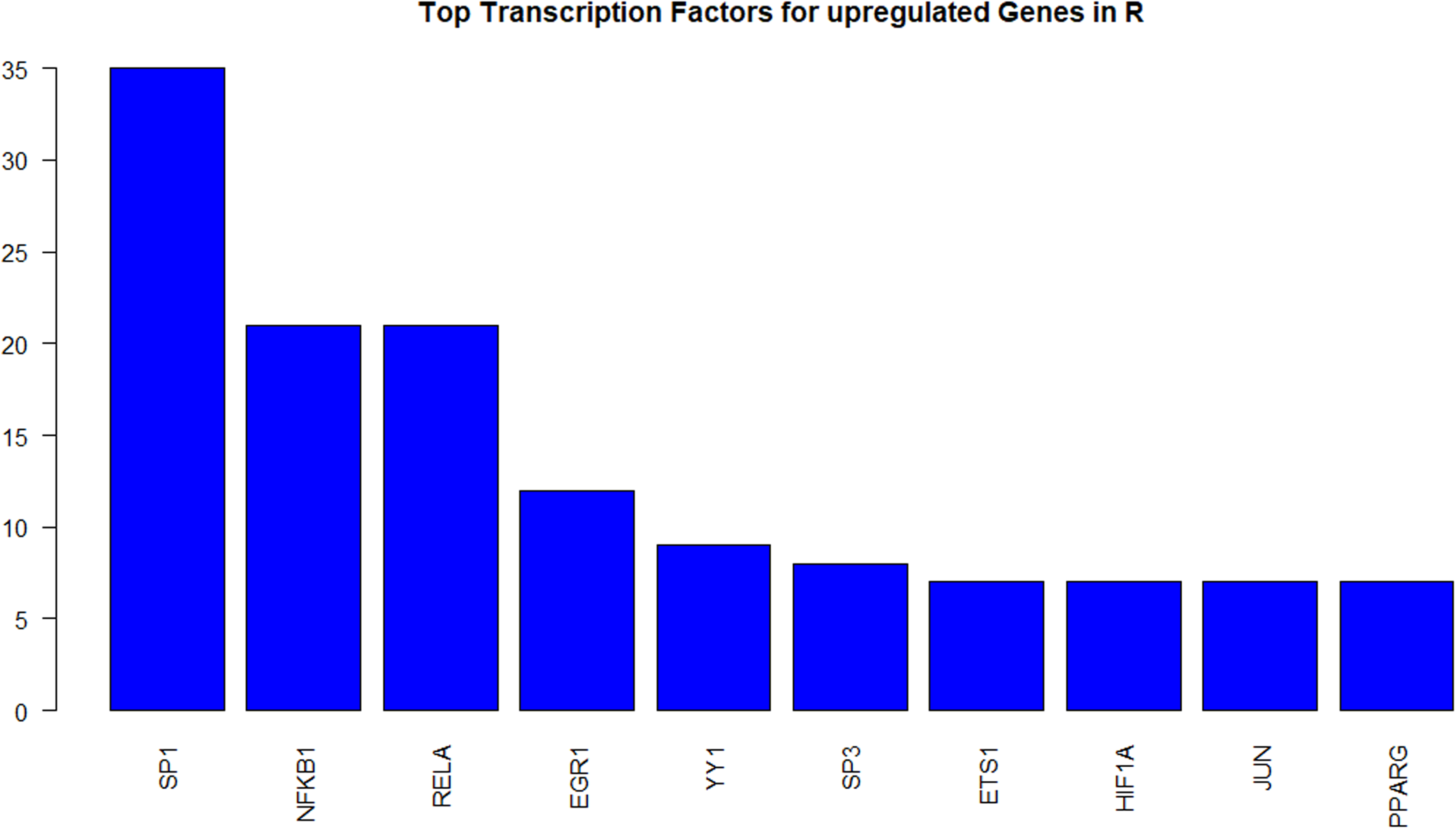

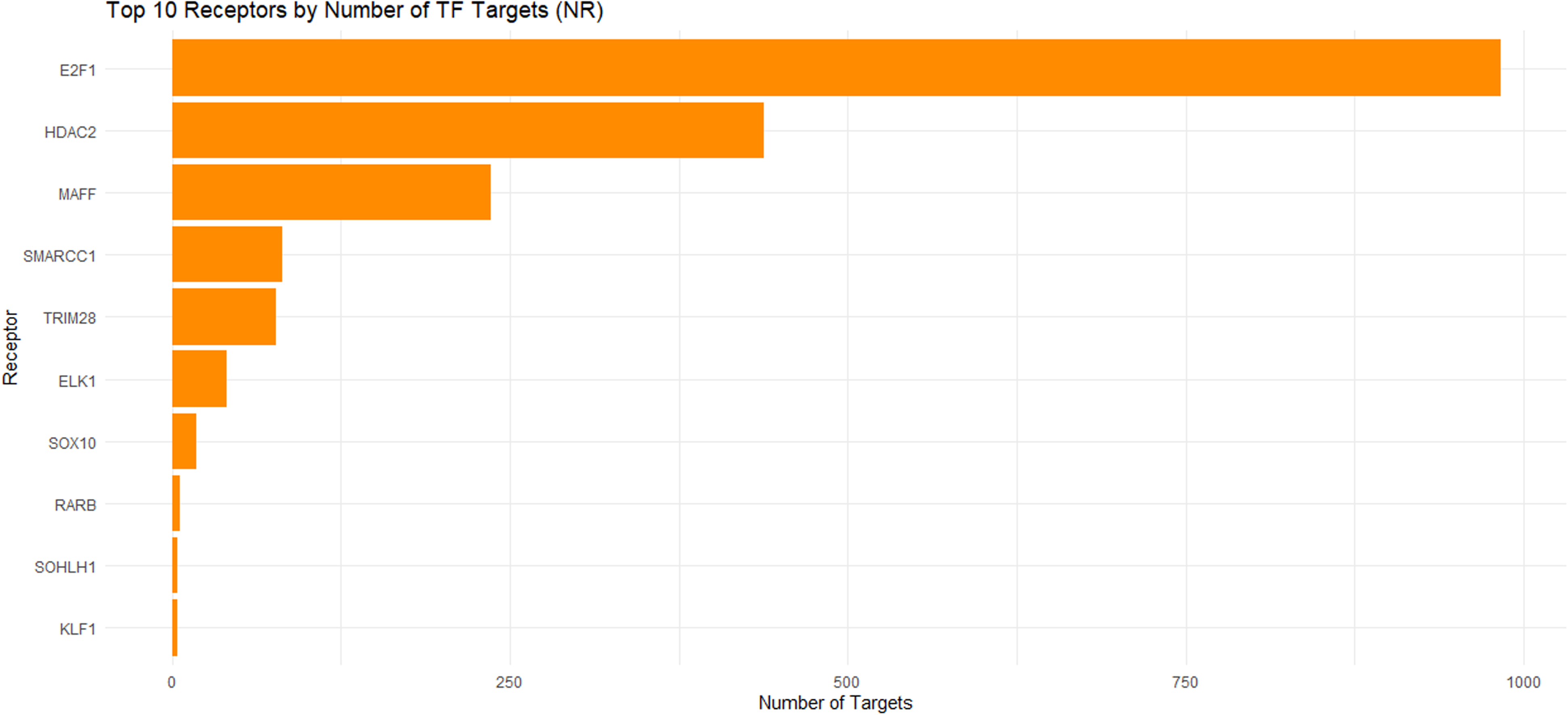

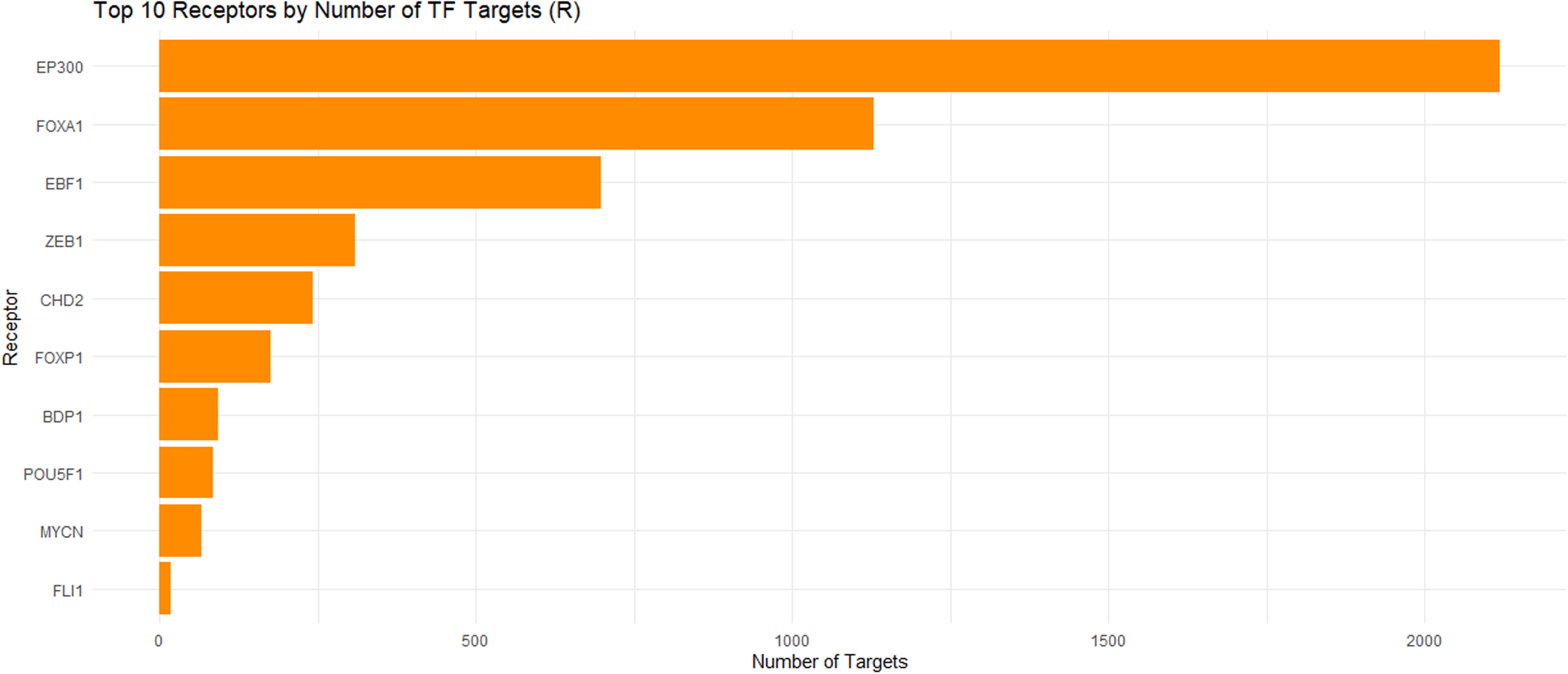

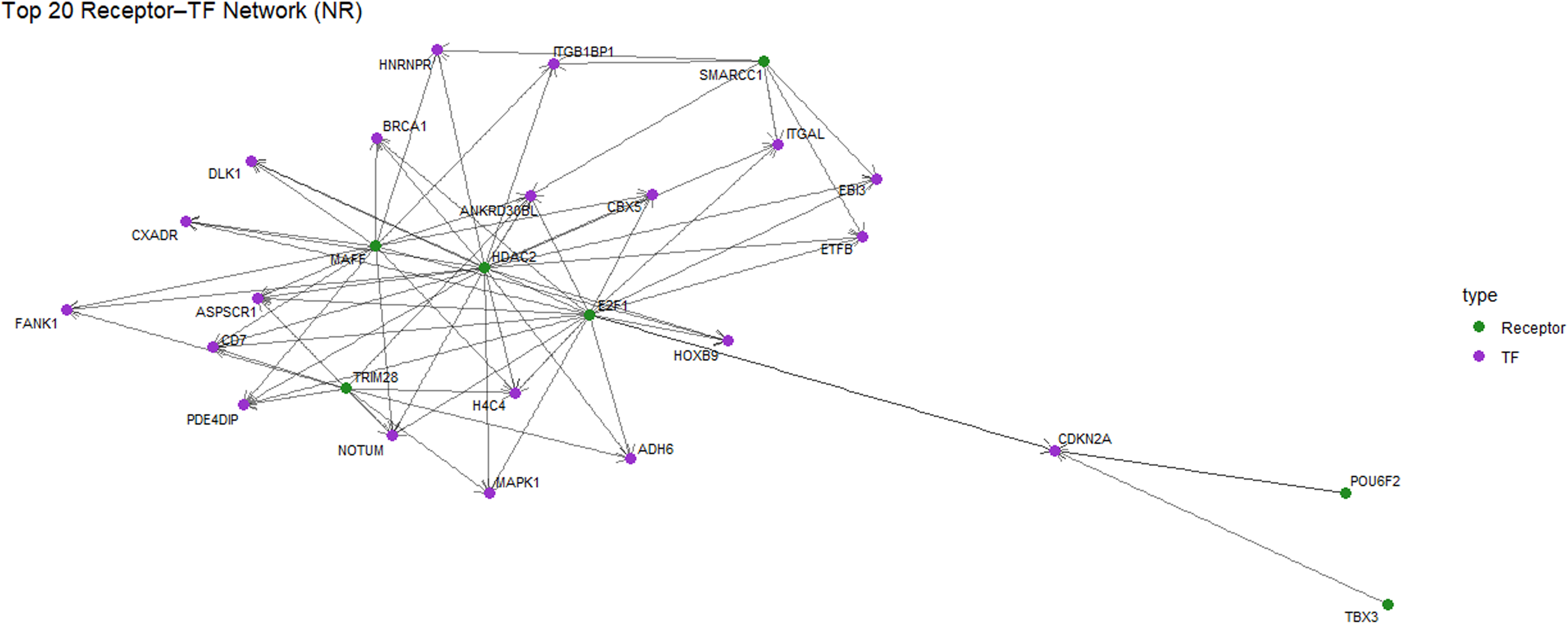

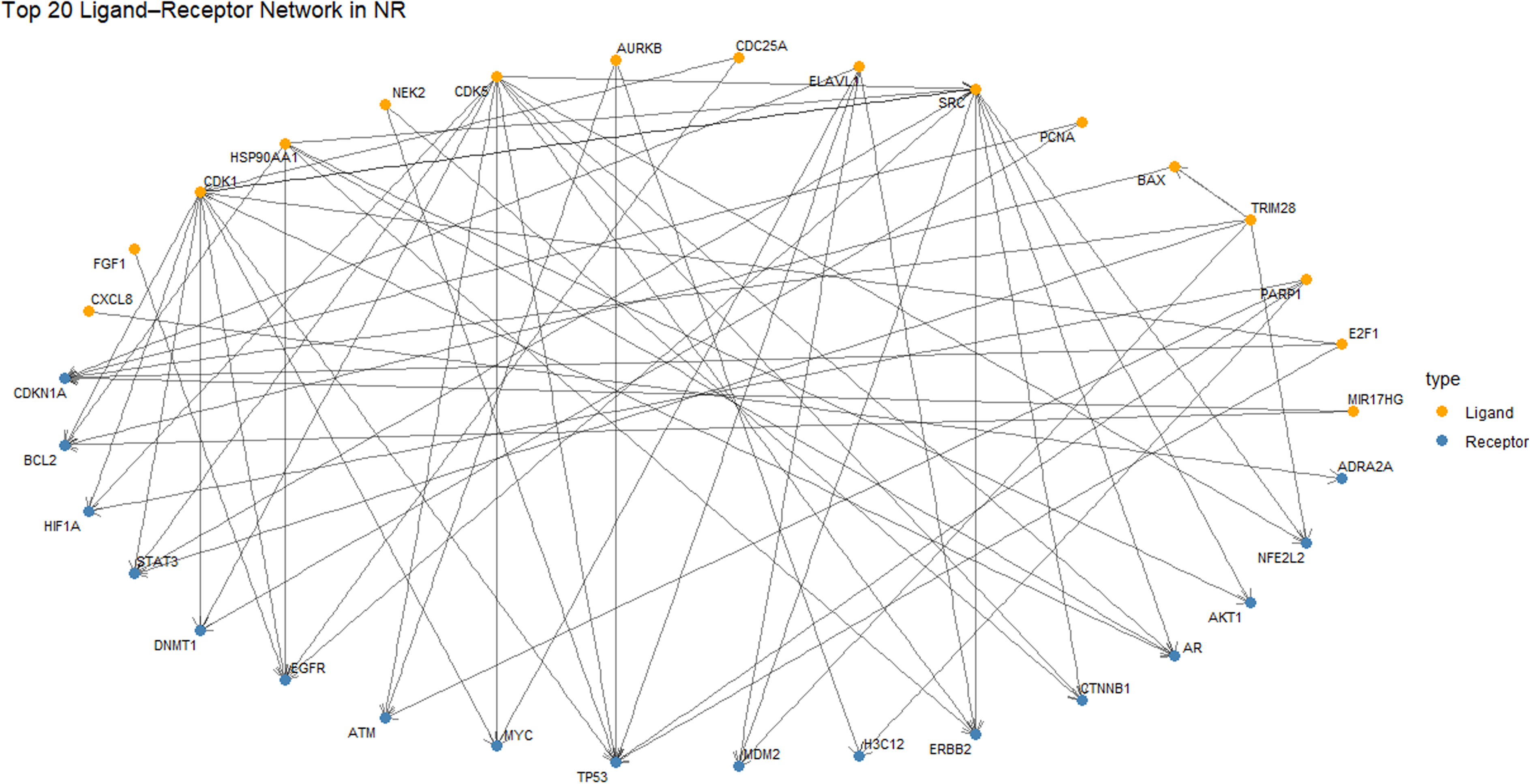

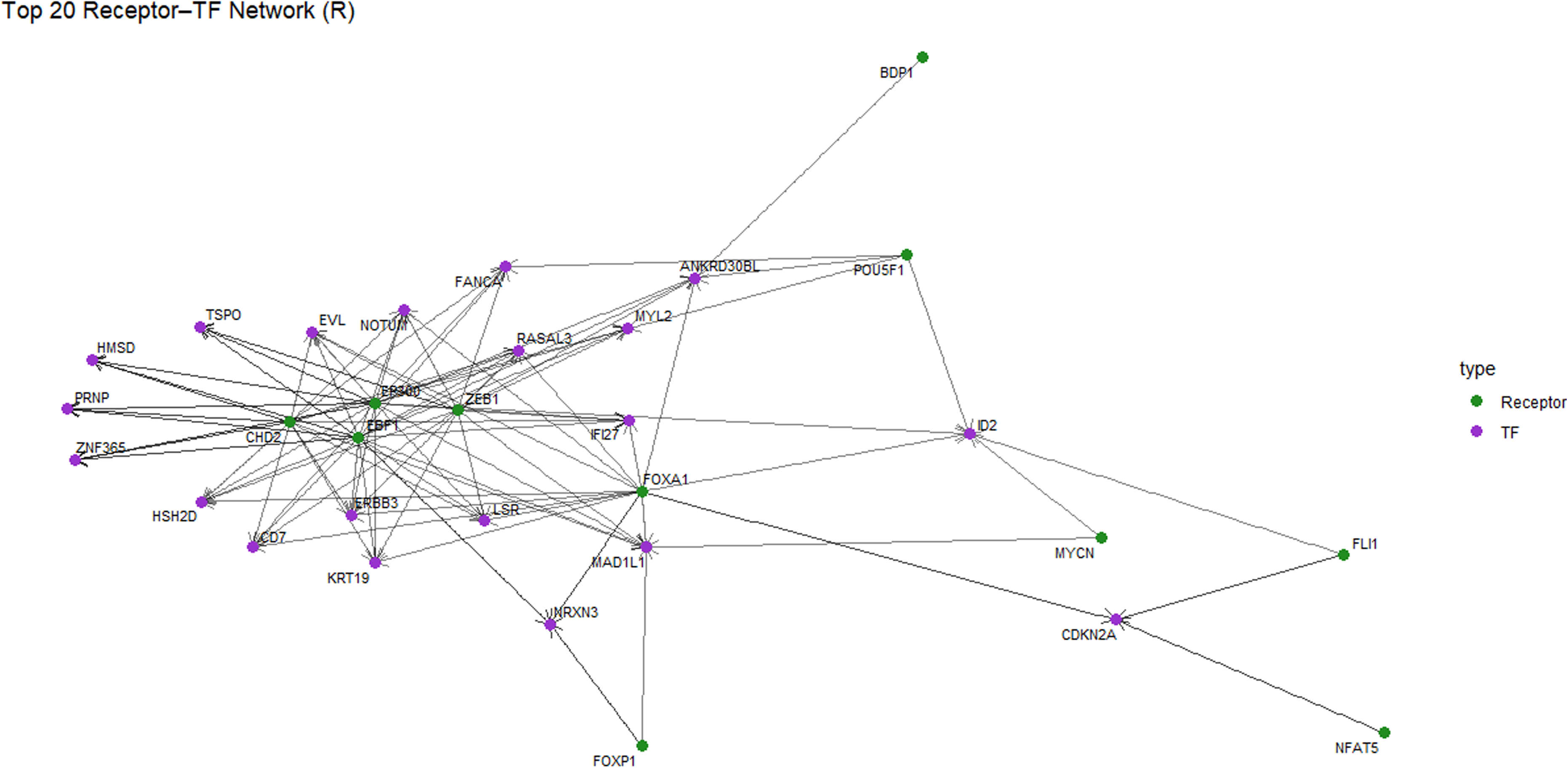

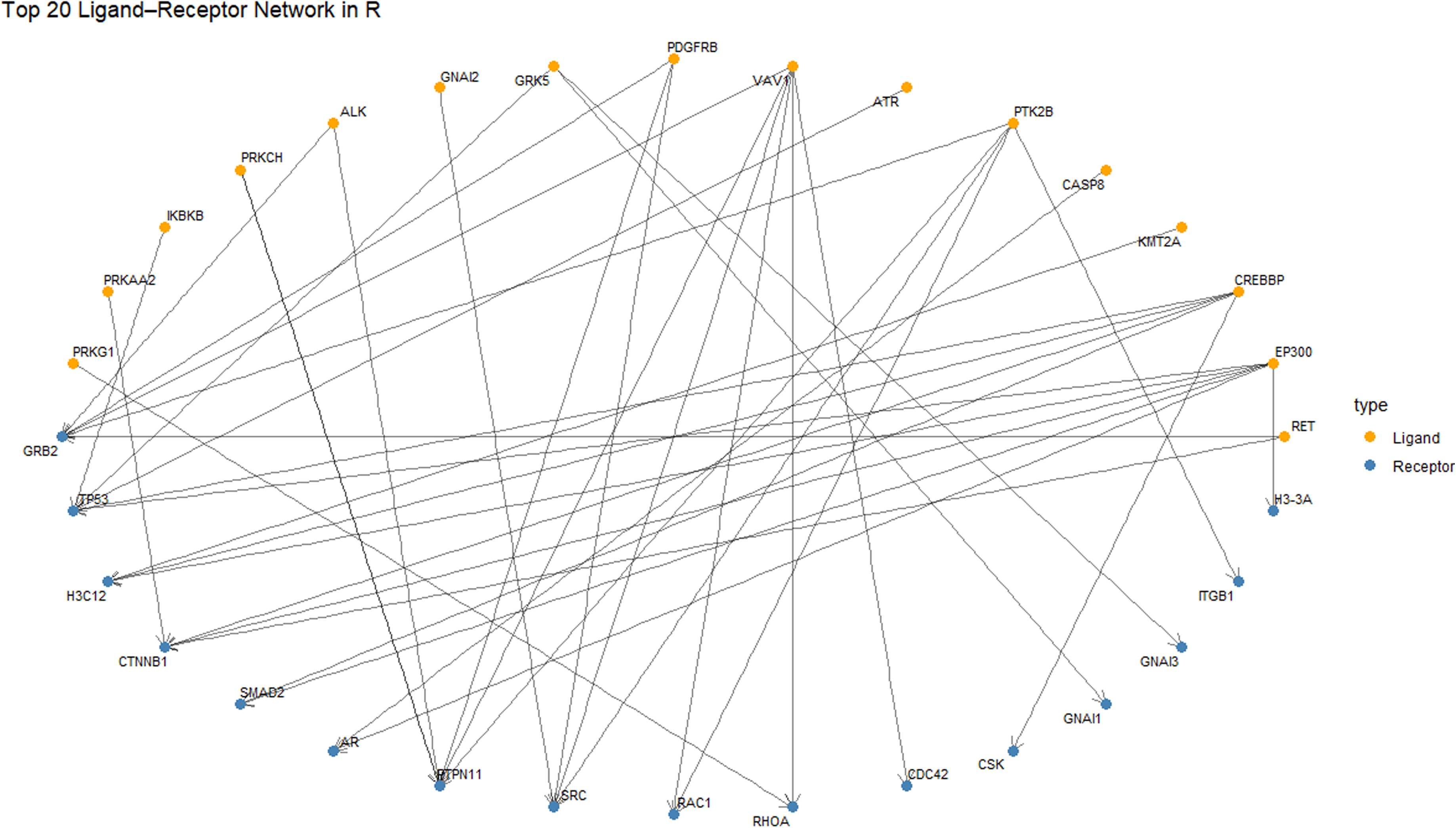

